# An atlas of spider development at single-cell resolution provides new insights into arthropod embryogenesis

**DOI:** 10.1101/2022.06.09.495456

**Authors:** Daniel J. Leite, Anna Schönauer, Grace Blakeley, Amber Harper, Helena Garcia-Castro, Luis Baudouin-Gonzalez, Ruixun Wang, Naïra Sarkis, Alexander Günther Nikola, Venkata Sai Poojitha Koka, Nathan J. Kenny, Natascha Turetzek, Matthias Pechmann, Jordi Solana, Alistair P. McGregor

## Abstract

Spiders are a diverse order of chelicerates that diverged from other arthropods over 500 million years ago. Research on spider embryogenesis, particular studies using the common house spider *Parasteatoda tepidariorum*, has made important contributions to understanding the evolution of animal development, including axis formation, segmentation, and patterning. However, we lack knowledge about the cells that build spider embryos, their gene expression profiles and fate. Single-cell transcriptomic analyses have been revolutionary in describing these complex landscapes of cellular genetics in a range of animals. Therefore, we carried out single-cell RNA sequencing of *P. tepidariorum* embryos at stages 7, 8 and 9, which encompass the establishment and patterning of the body plan, and initial differentiation of many tissues and organs. We identified 20 cell clusters, from 18.5k cells, which were marked by many developmental toolkit genes, as well as a plethora of genes not previously investigated. There were differences in the cell cycle transcriptional signatures, suggestive of different proliferation dynamics, which related to distinctions between endodermal and some mesodermal clusters, compared with ectodermal clusters. We found many Hox genes were markers of cell clusters, and Hox gene ohnologs often were present in different clusters. This provided additional evidence of sub- and/or neo-functionalisation of these important developmental genes after the whole genome duplication in the arachnopulmonate ancestor (spiders, scorpions, and allies). We also examined the spatial expression of marker genes for each cluster to generate a comprehensive cell atlas of these embryonic stages. This revealed new insights into the cellular basis and genetic regulation of head patterning, hematopoiesis, limb development, gut development, and posterior segmentation. This atlas will serve as a platform for future analysis of spider cell specification and fate, and studying the evolution of these processes among animals at cellular resolution.

## Introduction

Studying the embryology of arthropods, particularly insects, has helped to identify toolkit genes and their roles in development. It has elucidated ancestral mechanisms of developmental regulation on one hand, and how these processes evolve on the other [1]. Chelicerates, including spiders, represent an outgroup to mandibulate arthropods. Studying their development provides a unique perspective to better understand the evolution of embryogenesis among arthropods and other animals [2].

The common house spider *Parasteatoda tepidariorum* has proven to be a powerful model for understanding the genetic regulation of key processes during spider embryogenesis and specific spider innovations [2–6]. *P. tepidariorum* embryos initially form a radially symmetrical germ disc in one hemisphere, with the extra-embryonic and yolk tissue in the other [6–10]. Radial symmetry is broken during embryonic stages 5 and 6 to form a germ band by stage 7 [6, 11–13]. Therefore, generation of the bilaterally symmetrical germ band with both antero-posterior (A-P) and dorso-ventral (D-V) axes by stage 7 is a key point in embryogenesis. Subsequently, stages 7 to 9 encompass several important developmental events: *decapentaplegic*/*short gastrulation* mediate patterning along the D-V axes, and Hox instruct segment identity [8, 14, 15], the germ layers begin to differentiate into the corresponding[16] tissues and organs [16, 17], the head forms and neurogenesis begins [8, 18–20], the prosomal (cephalothorax) segments form [12, 21–23], concomitant with formation and growth of the limb buds [2, 6, 24] and the opisthosomal (abdominal) segments are added sequentially posteriorly from the segment addition zone (SAZ) [17, 25, 26]. Therefore, embryonic stages 7 to 9 see both linear changes in cell states as they differentiate into growing structures and tissues, and reiterative regulatory processes to generate the body along the A-P axis.

To better understand these processes we require differential gene expression data at cellular resolution during these stages of embryogenesis. Single-cell RNA sequencing (scRNA-seq) transcriptomics can provide this to allow cell state characterisation and differentiation and unbiased identification of key marker genes [24, 27–30]. Indeed, scRNA-seq has successfully been applied to embryos of a rapidly growing number of animals [31–38], including stage 5 and 6 *P. tepidariorum* embryos, which identified cells corresponding to the three germ layers and reconstructed A-P polarity and initial patterning based on known and new markers genes [39].

To better understand spider embryogenesis, we carried out scRNA-seq at stages 7, 8 and 9 stages taking advantage of our recent advances in cell dissociation, combining ACetic-MEthanol (ACME) dissociation [29] and SPLiT-seq scRNA-seq [40] technology. While most enzymatic dissociation methods process live cells, which can incur damage and stress related transcriptional signatures, ACME circumvents these problems by fixing cells during the dissociation process. The stages profiled are separated by just a few hours of developmental time, and ACME crucially fixes those stages avoiding hours of live cell dissociation where the cells may keep their developmental process *ex vivo*. Additionally, many droplet-based methods are sensitive to the introduction of noise from ambient RNA and cellular debris, which can also falsely increase their gene and UMI per cell counts. In contrast SPLiT-seq minimises these issues because ambient RNA is eliminated with the supernatant in each of the successive centrifugation steps for each of the split-pool rounds. Furthermore, the introduction of a FACS sorting step immediately prior to cell lysis eliminates cellular debris. Therefore, while SPLiT-seq usually results in lower gene and UMI per cell counts, compared to some other technologies, this approach resolves clusters that are robust to clustering conditions and subsampling.

Our analysis of stage 7, 8 and 9 spider embryos combining ACME dissociation and SPLiT-seq scRNA-seq allowed us to define cell types and capture new genes involved in several important developmental processes during these key embryonic stages, including germ layer differentiation, Hox patterning, head and CNS development, and limb development as well as new perspectives on the role of the so called ‘extra-embryonic’ cells. Furthermore, our results provided new insights into the regulation of the reiterative formation of the opisthosomal segments, most of which are sequentially generated from a posterior SAZ during stages 7, 8 and 9.

## Results

### Single-cell sequencing of three stages of spider embryogenesis

To better understand cell states during spider embryogenesis we sequenced single-cells from three embryonic stages (7, 8.1 and 9.1) of *P. tepidariorum* (Fig 1A) [6]. We focused on these stages because they mark the onset and/or continued progress of key developmental processes, including segmentation, and we lack information about the cells and their expression profiles during these processes [5, 6, 8, 11, 12, 14, 16–21, 25, 26, 41–50].

**Fig 1.**
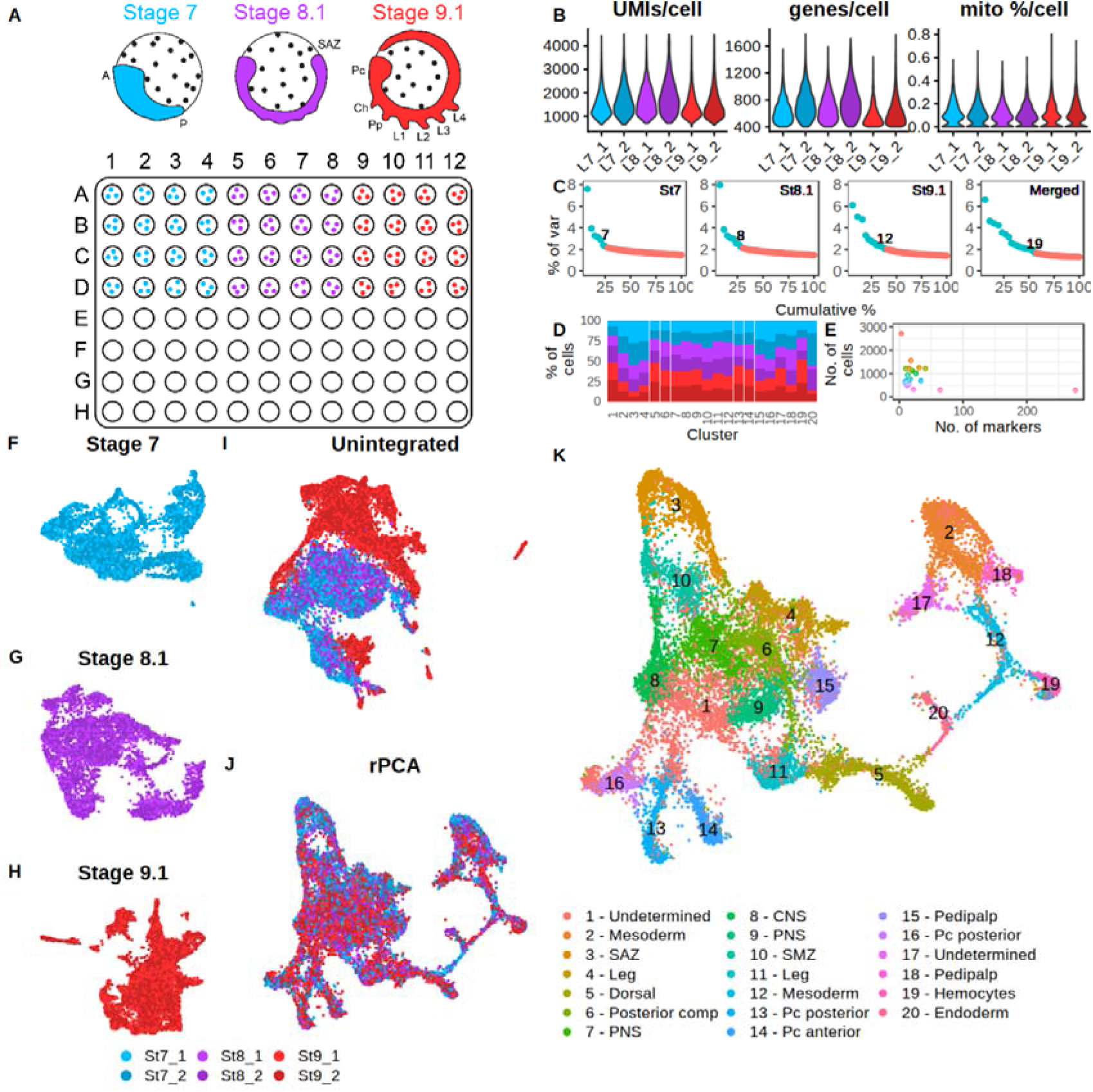
Single cell sequencing of three embryonic stages of *P. tepidariorum.* (**A**) Stages 7, 8.1 and 9.1 were collected, dissociated with ACME, and cells from each dissociation were plated as shown for the first round of SPLiT-seq barcoding. (**B**) Metrics of UMI, genes and mitochondrial expression in each library. (**C**) Significant PCAs per stage and all stages merged shows the significant PCAs increase from stage 7 to 9.1, with merged containing the most. (**D**) Percentage of cells from each stage/library for each cluster, normalised by taking an equal number of random cells from each stage. (**E**) Association between number of cells and markers per cluster. (**F** - **H**) UMAP for each stage. (**I**) UMAP for all stages merged without integration and (**J**) with rPCA integration. (**K**) UMAP and cell clustering with annotation derived from ISH of marker genes.

We carried out separate ACME dissociations for each stage [29]. These dissociation samples were subjected to SPLiT-seq barcoding [40], processing all stages in parallel (Fig 1A). Cells from each stage were barcoded separately so that first barcode could be used for de-multiplexing stages (Fig 1A). After the third round of SPLiT-seq barcoding, cells were FACS sorted into two samples for the fourth round of barcoding to generate two sequencing libraries. These libraries therefore constitute different cells from the same dissociation, that had different sequence indexes, and were pooled and sequenced on the same Illumina lane. A total of 481,741,227 raw sequence reads were generated from these two libraries (Sup Table 1). After trimming and barcode assignment using DropSeq tools, 328,915,611 (68%) reads remained (Sup Table 1). Star mapped 82% of input reads to a new *P. tepidariorum* annotation that was constructed to maximise 3’ completeness including UTRs to improve mapping (Sup Table 1). From these mapped genic reads, a total of approximately 30,000 cell transcriptomes were captured with a minimum of 100 genes, prior to Seurat filtering and doublet removal. Nearly all cells had <1% mitochondrial gene expression, indicating minimal transcriptional noise from cell stress in the dataset (Fig 1B) consistent with the use of ACME. After filtering (see Materials and Methods) based on UMI counts, genes expressed per cell, mitochondrial expression and doublet removal, the total processed dataset contained 18,516 cells, with 14,370 (43%) genes expressed out of a total 33,413 genes that we annotated in *P. tepidariorum*. Stages 7, 8.1, and 9.1 were represented by 4824, 4833, 8859 cells, respectively, with median UMI count per cell of 1465, 1656, and 1343, and a median of 674, 713, and 563 genes quantified per cell (Fig 1B & Sup Table 1). Stage 7 and 8 were comparable in these metrics, whereas stage 9.1 had fewer UMI and gene counts per cell, but more cells overall, which may reflect smaller cell sizes at stage 9 (Fig 1B & Sup Table 1).

Merging of cells from the two libraries for each stage and Seurat processing showed that libraries of each stage were comparable (Fig 1F - H). Cells from each library were distributed across UMAPs for each stage, with clusters containing cells from both libraries, suggesting no issues during the fourth round of barcoding and library preparation. Processing of each stage separately revealed an increase in the contribution of informative principal components increasing from 7 to 8 and 12 for embryonic stages 7, 8.1 and 9.1, respectively (Fig 1C). This suggests, as expected, that transcriptomic complexity increased as development progressed. Markers from each stage were identified using an in-cluster-versus-all-others, identifying 130, 117, and 230 markers for stages 7, 8.1 and 9.1, respectively. All markers are provided in Sup File 1.

### Stage sample integration

We assessed all stages together by merging and processing them without integration. This showed that stage 9.1 differed from stages 7 and 8.1 because there were clusters containing only/mostly stage 9.1 cells (Fig 1I). This suggested that: 1) there may be large differences between samples due to independent dissociations of embryos from each stage; 2) or that the greater number of cells from stage 9.1, and lower median UMI and gene counts, cause biologically similar cell states to appear transcriptionally different; 3) or that there are real biological transcriptional signatures at stage 9.1 causing cells not to be clustered with stage 7 and 8.1. For instance, the time interval between stages 7 and 8.1 is with 5-14 hours is much shorter than the 11-24 hours’ time between stage 8.1 and 9.1, which might explain the separation of stages [6].

We therefore assessed different integration approaches and explored their impact on cluster markers from the three stages by comparing the results to the unintegrated data. Since integration can force cell states to appear more comparable, information from stage 9.1 might be lost during the integration, and cause exclusion of stage 9.1 specific marker genes.

For integration we used both CCA and rPCA from Seurat, as well as Harmony [51]. We performed the same pre-processing, variable gene selection and normalisation prior to integration. Seurat rPCA, CCA and Harmony produced similar results, with only Harmony appearing to not integrate stage 9.1 as strongly as rPCA/CCA (Sup Fig 1). We therefore chose to proceed with rPCA since markers from both, the unintegrated and Harmony methods, were all also included in the rPCA marker list. In addition, rPCA is considered a more conservative approach that avoids over-correction [27, 52].

To establish whether integration generated data artefacts, or unlikely clustering patterns, we iterated through integration anchors (5 – 45), which affects the strength of integration, using a range of variable gene thresholds (1.2 – 1.7) (Sup Fig 2 & Sup Fig 3). Quantification of clustering similarity using an adjusted RandIndex [53] showed that the threshold of variable genes had the most effect on clustering similarity, and that integration with 30 to 40 anchors were most stable (Sup Fig 3). Therefore, we proceeded with rPCA integration with 1.3 threshold for variable genes and 40 number of anchors to assess all stages together.

To determine clusters in the integrated data we estimated a stable clustering resolution given the lack of information regarding spider cell type diversity. This approach revealed 20 clusters that were represented by cells from all stages/samples (see Materials and Methods) (Fig 1J & K), but still showed variability in the abundance of cells from a given stage within each cluster (Fig 1D), and cluster sizes ranged from 2719 (14.7%) cells in cluster 1, to 283 (1.5%) cells in cluster 20 (Fig 1E).

Marker genes for the 20 clusters were predicted with an in-cluster-versus-all-others approach, including genes that were expressed in at least 25% of cells in their respective cluster and an adjusted *p*-value return threshold of 1e-5. A total of 491 genes were identified as cluster markers, with numbers of markers per cluster ranging from 3 (cluster 1) to 276 (cluster 20) (Fig 1E). Cluster markers for integrated and stage data are provided in Sup File 1.

Given that the clustering showed sufficient structure and information we then interrogated cell clusters and characterised marker genes during these three stages with respect to key developmental processes. To assess whether stage-specific clusters were well-represented in the merged datasets we compared stage-specific cluster markers to unintegrated and integrated (rPCA) marker lists with a hypergeometric distribution test. Overall, we detected clear signatures that the unintegrated and rPCA integrated data possessed clusters that were also represented in stage-specific clusters (Sup Fig 4 & Sup Fig 5). While stages 7 and 8.1 had most of their markers present in the integrated marker list, stage 9.1 had several tens of markers missing. However, these missing markers were predominantly from a stage 9.1 cluster that related to cluster 19 of the integrated marker list both in terms of marker list overlap and in situ hybridisation expression of markers (Sup Fig 6). Therefore, integration helped to merge stages without considerable loss of stage-specific information.

### Clusters with the greatest G1 phase ratio relate to cells from gut, dorsal, and mesoderm lineages

Many new cells are required to build the differentiating germ layers, tissues, and organs during stages 7 to 9 of *P. tepidariorum* embryogenesis. We therefore asked whether any of the cell clusters were associated with signals of enhanced cell division and what tissues they may contribute to. We identified orthologs of *Drosophila melanogaster* genes that are associated with G1, S, G2/M phases of the cell cycle and quantified their expression in clusters using Seurat cell cycle scoring.

Sixteen clusters had similar proportions of each cell cycle phase, however the G1 phase was found in over 25% of cells in clusters 5, 12 and 19, and approximately 75% for cluster 20 (Fig 2A). This suggests that these four clusters exhibit different proliferation dynamics from other clusters, although none of the cell cycle genes were markers of these four clusters, or any other clusters. We next assessed the embryonic expression of markers from these four clusters.

**Fig 2.**
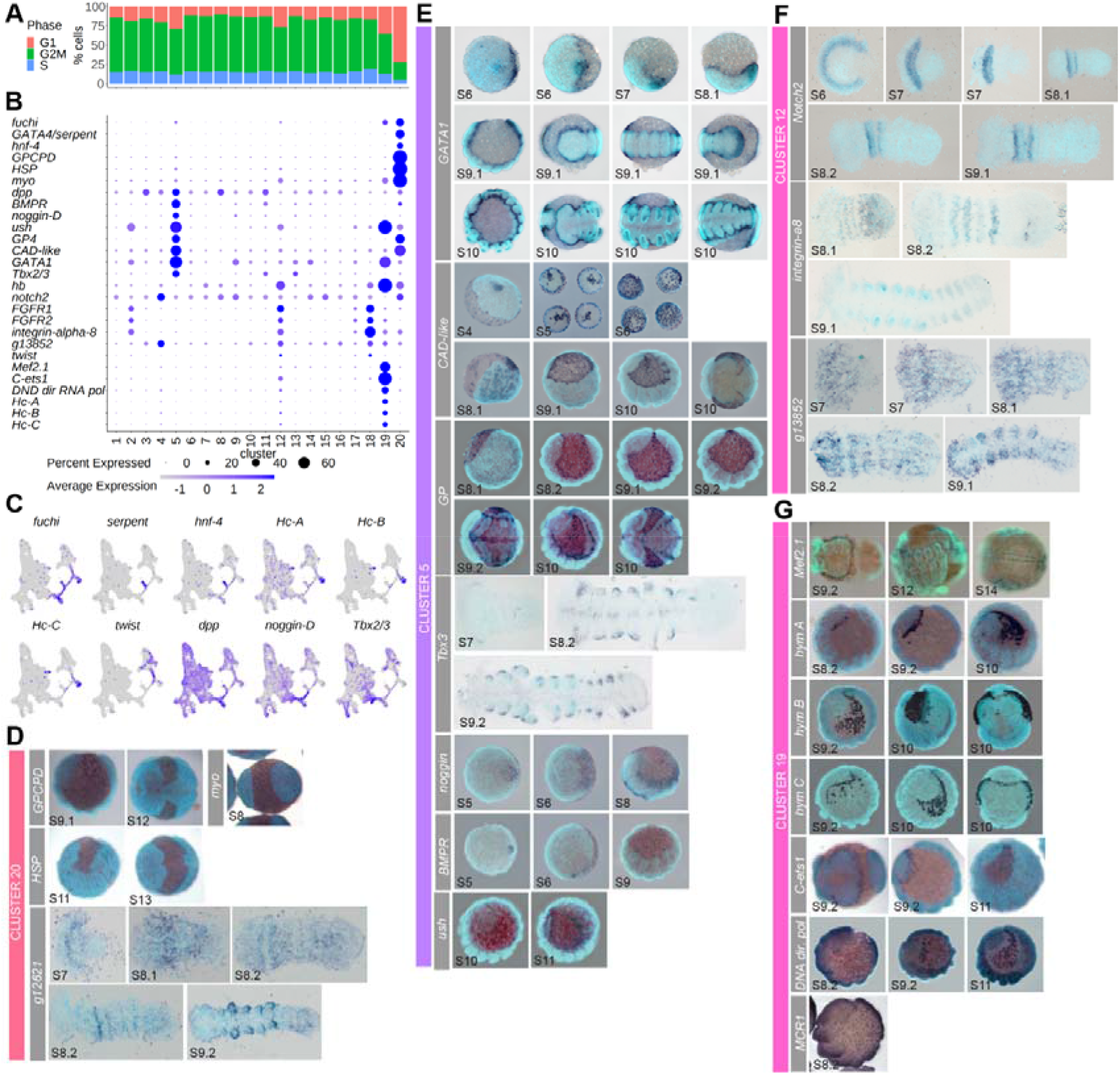
Cell cycle differences reveal four clusters with distinct endodermal and mesodermal characteristics. (**A**) Cell cycle scoring shows four clusters, 5, 12, 19 and 20, with more than 25% G1 phase cells (red). (**B**) Dotplot of marker genes. (**C**) UMAPs of some markers and non-markers previously identified to have similar expression as markers expressed in clusters 5, 12, 19 and 20. (**D** - **G**) Spatial expression of marker genes during embryogenesis assayed by in situ hybridisation.

Cluster 20 contained the highest number of significant marker genes (Fig 1E). We found that cluster 20 was marked by the recently characterised GATA gene *fuchi* [g16870], which is expressed in the endoderm [54]. These cells also expressed *hepatocyte-nuclear factor-4* (*hnf-4*) [g4057] and *serpent*/*GATA4* [g7067], which are also expressed in the endoderm, although they were not significant markers of cluster 20 (Fig 2B & 2C) [48, 54]. We analysed three further markers of cluster 20 *GPCPD* [g1958], *HSP* [g14898] and *myo* [g6459] (Fig 2B - 2D). Like *fuchi*, *serpent* and *hnf-4*, all three marker genes were expressed in extra-embryonic cells that contribute to the endoderm (Fig 2D). *GPCPD* was previously used as an endoderm marker, and this gene is expressed earlier in the peripheral cells of the germ-disc and in the cumulus [12] (a group of cells that migrates during stage 5 to determine the dorsal field at stage 6 [8, 9, 12]), further evidencing that this cluster relates to endoderm cells.

Cluster 5 was marked by the dorsal determinant *dpp* [g29377] [8], as well as *BMPR* [aug3.g2323], *noggin-D* [g27229], *ush* [aug3.g16893], *platelet glycoprotein 4* (*GP4*) [g4985], *CAD-like* [g6385], *GATA1* [g4744] [54] and *Tbx2/3* [aug3.g3745] (Fig 2B, 2C & 2E). Like *dpp*, genes such as *BMPR*, *CAD-like, noggin-D* and *GATA1* were all expressed in the cumulus, indicating that cluster 5 cells might originate from the mesenchymal cumulus cells underneath the epithelial cells and have dorsal identity [8, 9, 23, 39]. Indeed, several of these markers (*noggin-D*, *GATA1* and *BMPR*) were present in the cumulus cell cluster in a recent single-cell analysis of stage 5 embryos [39]. We found that *GATA1*, *BMPR, ush*, *GP4* and *noggin-D* were all expressed in the dorsal field at stage 6 (Fig 2E). *noggin-D* and *GATA1* were also expressed from stages 7 to 9 broadly around the dorsal periphery of the germ band. Previous analysis showed that the embryonic expression of *GATA1* borders ventral *short gastrulation* (*sog*) expression at stage 8 [12]. However, *BMPR* expression was restricted to dorsal domains in each appendage at stages 7 to 9 (Fig 2E). *Tbx2/3* was expressed in the precheliceral region and at the ventral midline, as well as in dorsal regions of prosomal appendages, like *BMPR* (Fig 2E). We observed that in addition to the expression of *CAD-like* in the cumulus, there was also expression in large cells surrounding the germ disc and in the extra-embryonic region at stage 5 (Fig 2E). Subsequently, *CAD-like* expressing cells at stage 6 were present in the dorsal field and extra-embryonic region and at stages 7 to 8 beneath the germ band. At stage 9 expression of this gene is present in extra-embryonic cells around the germ-band, not underneath the germ-band, like all other cluster 5 markers surveyed (Fig 2E). These results suggest that cluster 5 relates to cells originating from the cumulus that form the dorsal region of the germ band and contains cells located in the extra-embryonic region.

Cluster 12 was marked by *hunchback* (*hb*) [g27583], which was previously shown to be necessary for development of the L1, L2 and L4 prosomal leg-bearing segments (Fig 2B) [43]. Another cluster 12 marker, *Notch2* [g30344], was first expressed in a single broad domain in the anterior of the germ disc at stage 6 that splits during stage 7 to 8 and, like *hb*, was subsequently expressed in L1 and L2 from stage 8 (Fig 2F) [43]. Cluster 12 was also marked by the mesodermal genes, *FGFR1* [g11749] and *FGFR2* [g3961] [13], as well as *integrin-alpha-8* [g23098] and an uncharacterised gene g13852, which were also expressed throughout the mesoderm of the prosomal and opisthosomal (Fig 2F). In a previous study *integrin-alpha-8* was also identified as a marker for a cluster at stage 5 and was expressed in cells that form a mesodermal cell lineage at the germ-disc periphery [39]. While *twist* [g22789], a mesodermal related gene in *P. tepidariorum*, was not a marker of cluster 12, it was expressed in cells of clusters 2, 12, 17, and 18 (Fig 2C). This suggests that cluster 12 represents mesoderm that originates from cells at the germ-disc periphery but later become broadly distributed across the germ-band.

Cluster 19 was marked by the mesodermal gene *Mef2.1* [g3542] (Fig 2B & 2G) [55, 56]. *Mef2.1* and three other markers, *hemocyanin A* (*Hc-A*) [g11873], *C-ets1* [g472], and *DNA directed RNA pol* [g17128], all showed expression from the dorsal regions around the head into the extra-embryonic region (Fig 2G). Two other hemocyanin genes (*hemocyanin B* [g13621] and *hemocyanin C* [g22680]) that were markers of stage 9.1, cluster 11, which had the best marker overlap with cluster 5 from the integrated data (Sup Fig 5), had similar expression to *hemocyanin A* (Fig 2G). The post stage 9 expression of these marker genes suggests either dorsal cells are migrating across the extra-embryonic region earlier than dorsal closure at stage 13 [6], or that extra-embryonic cells are being recruited to dorsal tissues of the embryo proper. Collectively, given the function of the orthologous genes in *D. melanogaster* [57–60], cluster 19 cells likely correspond to hemocytes, which until now have not been identified in spiders.

Overall, we were able to characterise endodermal and mesodermal cell populations, including a new identified hemocyte related population. Our dataset will serve as a resource for further characterising these cell populations during embryogenesis.

### Hox markers are consistent with A-P cell identity and evidence sub- and/or neo-functionalisation

*P. tepidariorum* has retained two Hox gene clusters following the whole genome duplication (WGD) event in an ancestor of arachnopulmonate arachnids and is missing only a second copy of *fushi tarazu* (*ftz*) [14]. During stages 7 to 9, the Hox genes are generally expressed in a collinear fashion across the A-P axis of *P. tepidariorum* embryos [14] and therefore we assessed the expression of these key patterning genes in the scRNA-seq data.

Expression of all nineteen Hox genes in *P. tepidariorum* was detected in the scRNA-seq data and ten were markers of eight clusters (clusters 2, 3, 4, 9, 10, 11, 15, 18) of the integrated data (Fig 3A & Sup Fig 7A & 7B). Fitting with the collinear expression principle, the posterior Hox genes were expressed at stage 9.1 more so than stages 7 and 8.1 (Fig 3B). Hierarchical clustering of our single cell data using Hox marker expression identified six groups (Fig 2C). Four of these corresponded to spatial regions of the spider embryo. Whereas the others potentially represent one group of spatially distributed cells expressing but not marked by Hox genes and another group of non-Hox expressing cells (Fig 2C) [14, 15].

**Fig 3.**
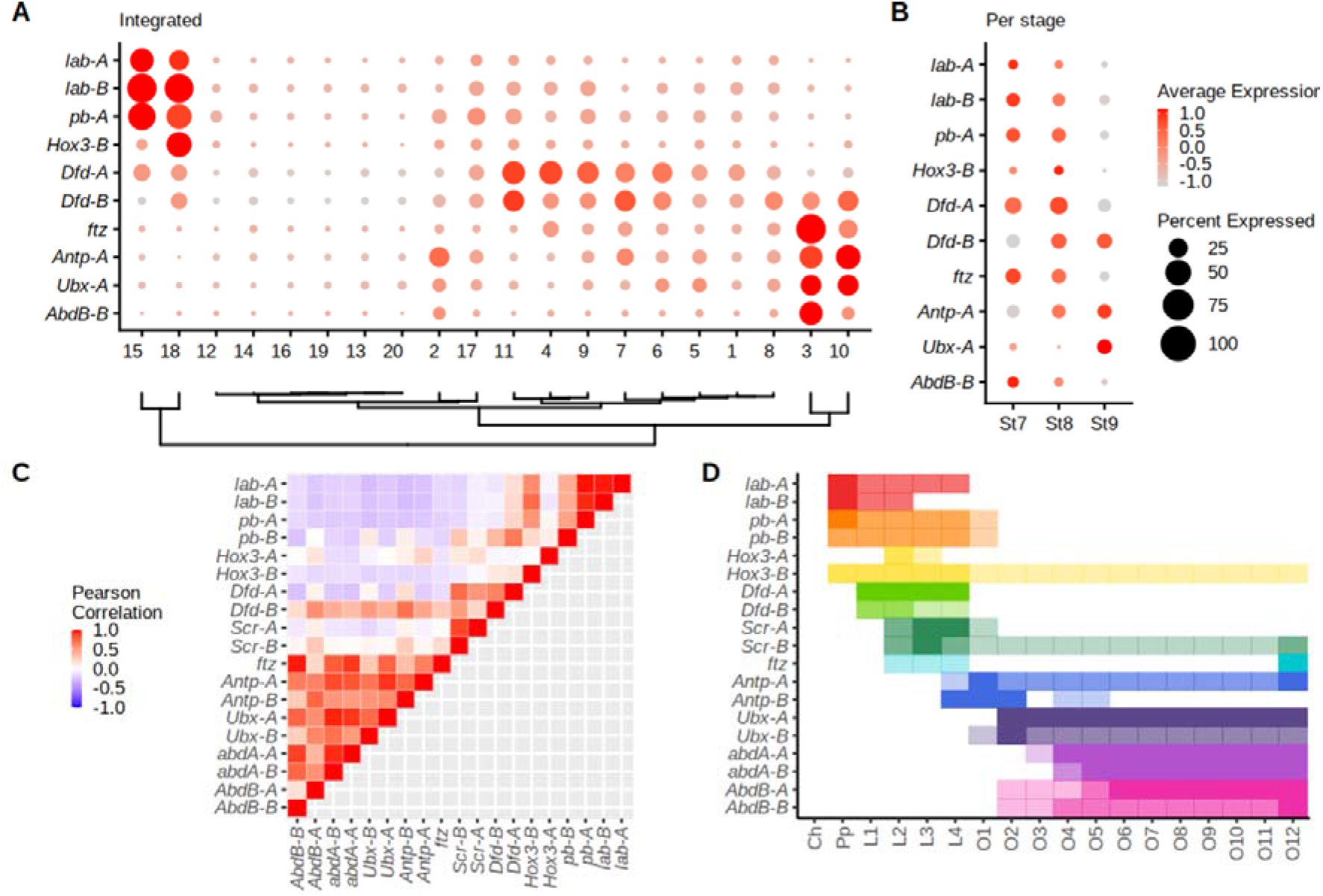
Hox expression in scRNA-seq data. (**A**) Dotplot of all ten Hox marker expression in cell clusters ordered hierarchically for integrated data. (**B**) Temporal expression of thirteen Hox markers across the three stages. The legend in (**B**) relates to the expression and number of expressing cells per cluster in both (**A)** and (**B)**. (**C**) Pearson’s correlation coefficients of SCT normalised average Hox expression across all 20 clusters reveal three cluster types related to pedipalps, legs and opisthosoma identities. (**D**) Positioning of Hox expression across the A-P axis of *P. tepidariorum*. Spatial expression intensity (detected by RNA in situ hybridisation) reflected using opaque (strong expression) to more transparent colouring (weak expression). Adapted from Schwager *et al* 2017, with the addition of new *Hox3-A* expression data (Sup Fig 7D).

The four groups that relate to spatially confined Hox expression suggest that some clusters represent cells that are restricted to segments across the A-P axis. One group related to the pedipalpal segment and contained clusters 15 and 18, which are marked by *labial-A, labial-B,* and *proboscipedia-A*, which are most highly expressed in the pedipalpal region (Fig 3C) [14, 15]. Another group relating to prosomal segments contained clusters 4, 9 and 11, marked by *Deformed-A* and *Deformed-B*, which are exclusively expressed in the leg-bearing segments (Fig 3C) [14]. Additionally, other markers of cluster 11 included *Distal-less* (*Dll*) [g10793] [44] and *sp6-9* [g22966], and a previously unanalysed marker *basonuclin* [g29744], which are also expressed in leg-bearing appendages (Sup Fig 7C) [61, 62], further supporting that these clusters contain cells that are found in prosomal and leg-bearing segments. The other two groups relate to the opisthosomal region, with one group containing clusters 3 and 10, marked by *ftz*, *Antennapedia-A*, *Ultrabithorax-A* and *Abdominal-B-B*, which are all expressed in the opisthosomal region, and another group containing cluster 2, marked only by one Hox gene, *Antennapedia-A* (Fig 3C) [14].

The remaining two groups comprised of clusters that did not have Hox markers. One of them, which contained clusters 1, 5, 6, 7 and 8, showed expression of many Hox genes but none passed statistical thresholds to become markers, suggesting that these cell clusters represent cells that are distributed across the A-P axis (Fig 3C). The other group contained clusters 12, 13, 14, 16, 19, and 20, which in contrast did not express any Hox gene, suggesting that there are populations of cells in *P. tepidariorum* embryos that are not patterned by Hox genes at stages 7 to 9.1 (Fig 3C).

We next assessed the correlation of Hox expression across clusters to see if ohnologs show similar or divergent expression across clusters (Fig 3D). Generally, this supported the hierarchical clustering showing distinct correlation groups relating to pedipalpal, leg-bearing and opisthosoma identities (Fig 3D). It also provided further evidence for the hypothesis that some Hox ohnologs have undergone sub- and potentially neo-functionalisation previously indicated by comparing spatial and temporal expression pattern analysis. *Hox3* ohnologs, for example have low correlation (r = -0.007), whereas other Hox ohnologs have highly correlated expression across clusters e.g *labial* (r = 0.97). Another example is the two *proboscipedia* (*pb*) ohnologs, which are both expressed in appendage mesoderm, but only *pb-A* is expressed strongly in the pedipalp segment [14]. This pattern is also reflected in the scRNA-seq data, whereby *pb-A* is a marker (of clusters 15 and 18), whereas *pb-B* is not a marker of any cluster and only spatially expressed lowly in several prosomal regions. Furthermore, *Scr-A* is expressed mostly in leg bearing segments, whereas *Scr-B* is also expressed in the SAZ [14] consistent with expression detected in cells with opisthosomal identity in the scRNA-seq data (Sup Fig 7B).

Overall, our analysis of Hox gene expression showed that the scRNA-seq data captured key developmental transcription factors and allowed us to compartmentalise many cell clusters into broad regions of the body plan, as well as clusters that were mostly void of Hox expression. Furthermore, this single cell data supports previous expression pattern analysis that spider Hox ohnologs have undergone sub- and potentially neo-functionalisation after WGD [14] and offers a resource for future investigations of other ohnologs.

### Patterning of the spider precheliceral region

The *P. tepidariorum* germ disc periphery represents the future anterior of the germ band that later gives rise to the precheliceral region as well as structures like the brain, mouth parts and eyes [18, 19, 45]. *hedgehog* (*hh*) and *orthodenticle-1* (*otd-1*) are co-expressed at the germ-disc periphery at stage 5 and begin to move posteriorly at stage 6 (Fig 4A) [15, 22, 45]. The most posterior domain of *hh*, which does not co-express *otd-1*, undergoes wave-splitting from stage 7 to 8 to first generate the *lab-A* expressing pedipalpal segment, and then the cheliceral segment (Fig 4A) [15, 22, 45]. However, the region of *hh* that co-expresses *otd-1* denotes the boundary of this *hh* wave-splitting process, and therefore represents the posterior boundary of the precheliceral region (Fig 4A). While *hh* and *otd-1* are involved in setting up the precheliceral region it is unclear how the precheliceral structures are thereafter patterned [45]. By assessing new and existing gene expression studies of cluster markers and per-stage-per-cluster markers, we identified three cell clusters, 13, 14 and 16, with different gene expression regionalisation and dynamics in the precheliceral region. These three clusters were marked by known anteriorly expressed genes, *otd-1* and *hh*, but lacked Hox markers (Fig 2, 4B & 4C), matching previous observations that the precheliceral region does not express Hox genes after the pedipalpal and cheliceral segments have been defined.

**Fig 4.**
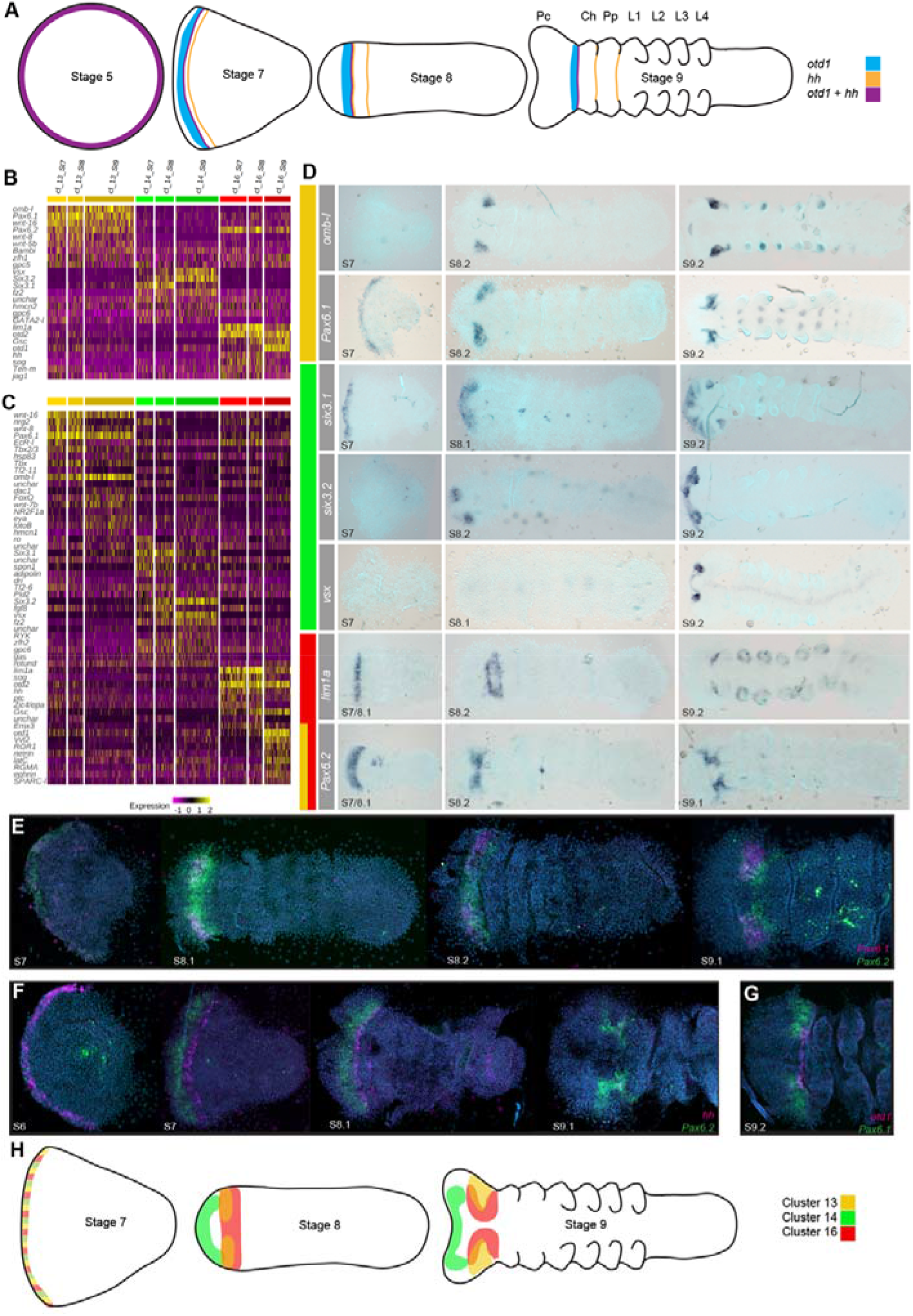
Clusters contributing to the spider precheliceral region. (A) Schematic overview of *otd-1* and splitting *hh* expression in prosomal patterning. At stage 5 *otd-1* and *hh* expression overlaps. At stage 7 expression of both genes migrates posteriorly from the anterior rim, and *hh* splits to form the presumptive pedipalp segment. At stage 8 the second splitting wave of *hh* forms the presumptive cheliceral segment. At stage 9 *otd-1* and *hh* are expressed at the posterior region of the precheliceral segment [22, 45]. (B) Heatmap of top ten markers for clusters 13, 14 and 16 relating to precheliceral region patterning. (**C**) Heatmap of top ten markers of the same clusters but also comparing between stages. These show some staggering of markers across stages, suggesting possible differentiation pathways (**D**) Expression of marker genes, colour bars represent the clusters they are associated with relative to figures B and C. (**E** – **G**) Double fluorescent in situ hybridisation of markers. (**E**) *Pax6.1* and *Pax6.2* markers relate to cells from clusters 13 and 16, which show expression moving from the anterior rim towards the posterior of the precheliceral region. *Pax6.1* expression is more dorsal and anterior compared to *Pax6.2*, with regions of non-overlapping expression (**F**). The most posterior expression of *Pax6.2* co-expresses *hh* (**H**) Overview of the location of proposed three head patterning clusters in the scRNA-seq data from stage 7 to 9.

o*td1* [g5047] and *hh* [g23071] were markers of cluster 16 (Fig 4B & 4C), suggesting this cluster relates to cells of the precheliceral region. We therefore analysed two other markers from cluster 16 (Fig 4B) and found that *lim1a* [g12191] and *Pax6.2* [g12868] were both expressed at stage 7 several cells posterior to the very anterior rim of the germ band (Fig 4D). Like *otd-1* the expression of *lim1a* and *Pax6.2* subsequently migrates more posteriorly, resulting in a broad band at the posterior boundary of the precheliceral region at stage 8 (Fig 4D). By stage 9 this band becomes restricted to two domains lateral to the ventral midline (Fig 4D). This is consistent with another cluster 16 marker *sog* [g13327], which is expressed in the ventral midline [8] and is expressed in stage 7 and 8 cluster 16 cells but has reduced expression by stage 9 (Fig 4C).

We assessed the cluster 13 markers *Pax6.1* [g12873], an *optomotor-blind* like gene [aug3.g3790], and *Tbx2/3*, which was also a cluster 5 marker (note that *Pax6.2* was also a cluster 13 marker – see above). *Pax6.1* and *Tbx2/3* were expressed earlier than the other cluster 13 markers. At stage 7, *Pax6.1* was expressed in a stripe along the anterior rim, and *Tbx2/3* had faint expression at the lateral edges of the germ band close to the anterior rim (Fig 4D). Their expression subsequently migrated towards the posterior of the precheliceral region (Fig 4D). However, in contrast to cluster 16 genes, most markers became limited to the posterior dorsal region of the precheliceral region by stage 9 (Fig 4D).

Cluster 14 cells were not marked by *otd-1* expression, however we analysed three markers, *six3.1* [g1245], *six3.2* [g25543], and *visual system homeobox (vsx)* [aug3.g27186], which were all expressed in the precheliceral region (Fig 4D). *six3.1* was initially expressed along the anterior rim of the germ band at stage 7, while *six3.2* appeared at the anterior rim at stage 8, and finally *vsx* by stage 9 (Fig 4D). All three genes (*six3.1*, *six3.2* and *vsx*) maintained their expression at the anterior of the precheliceral region (Fig 4D).

Double fluorescent in situ hybridisation of precheliceral region markers was performed to better distinguish the expression dynamics and mutually exclusive regions between clusters. *Pax6.1* (cluster 13) was initially maintained at the anterior limit of the *Pax6.2* (cluster 13 and 16) expression domain (Fig 4E). By stage 9.1 *Pax6.2* was more ventrally restricted and *Pax6.1* was more dorsally restricted, overlapping by a few cells, corroborating their distinct D-V domains of expression in the developing precheliceral region detected by single stainings (Fig 4E). *Pax6.2* remained anterior to the *hh* stripe from stage 7 to 8.1 and at stage 9.1 their expression overlapped at the posterior border of the precheliceral region (Fig 4F). This was further supported posterior *Pax6.1* expression overlapping with *otd-1* (Fig 4G).

Overall, our data revealed new details of transcription factor expression in the developing precheliceral region. It suggested that distinct combinatorial expression domains mark different regions of the precheliceral region such as the forebrain, and hindbrain primordia (clusters 13 and 16), which become more obviously regionalised during stages 8 and 9 (Fig 4H) [63, 64]. However, we appreciated that it is difficult to spatially place these cells across time from stage 7, since it is unknown whether gene expression is dynamic across cell fields, or if cell movement/recruitment underlie spatial changes in gene expression. Identification of markers between stages of each cluster in a per-stage-per-cluster approach reveals that all three clusters have transcriptional transitions between stages, suggesting there is dynamic expression (Fig 4H). Future work will reveal how these expression dynamics relate to differentiation gene regulatory networks in precheliceral patterning. Additionally, clusters 13, 14 and 16 had better overlap with clusters from stage 8.1 and 9.1 specific clustering, rather than with stage 7, consistent with the precheliceral region being established after stage 7.

### Ventral midline patterning and diversity in peripheral nervous system cells

Following D-V axis formation during stages 5 and 6 [8, 12], patterning of the ventral midline during stages 7 to 9 is critical for the development of the nervous system. Therefore, we next explored our single cell data to gain deeper insights on this process. Ventral patterning in *P. tepidariorum* is regulated by expression of *sog* [g13327] [8]. The strongest expression and marker association (adjusted *p*-value) of *sog* is present in cluster 8, although it was also a marker of clusters 4, 7, 10 and 16 (Fig 5B). We therefore assayed the spatial expression of five additional cluster 8 marker genes (*Nkx6.2* [g12201], *RGMA* [g28941], *LRR2* [g7463], *vitK-C* [g11868] and *hamlet* [aug3.g11431]). While the onset of their expression varied, they were all, like *sog*, expressed along the ventral midline and excluded from the posterior SAZ as well as the peripheral cells of the germ band by stage 9 (Fig 5A). Therefore cluster 8 is related to the ventral midline. This suggests that while there are multiple *sog* related ventral clusters, perhaps due to the broad expression of *sog* at stage 7 and 8.1 [8], cluster 8 was composed of cells that likely comprise the nerve cord cells, regulated by genes like *Nkx6.2*, which has a conserved role in specifying the ventral midline [65].

**Fig 5:**
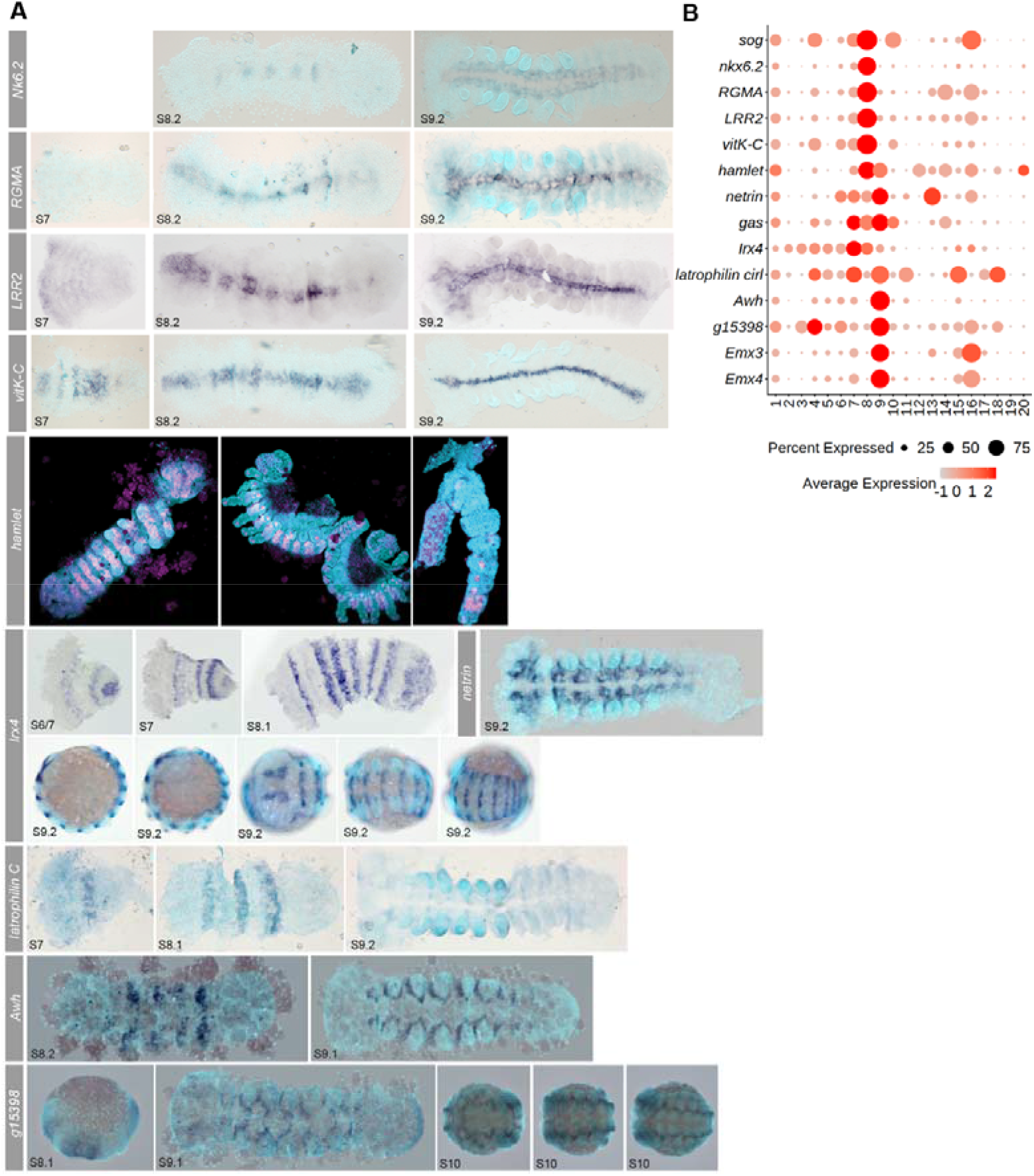
CNS and PNS related clusters in *P. tepidariorum*. (**A**) Single RNA in situ hybridisations of marker genes for clusters 7, 8 and 9. Cluster 8 markers *Nk6.2*, *RGMA*, *LRR2 vitK-C* and *hamlet* are all expressed through the ventral midline. Clusters 7 and 9 markers show metameric patterns of expression around the ventral midline and appendages, which relate to putative PNS cells. (**B**) Dotplot for markers of clusters that represent CNS and PNS cells.

Clusters 7 and 9 share the markers *hamlet*, *netrin* [g18008] (Fig 5A & 5B) and *growth arrested specification* (*gas*) [g1575] [50], which incidentally were also the only markers of cluster 1. All three of these genes had complex and broad expression across the embryo (Fig 5A). Other cluster 7 markers *Irx4* [g29290] [46] and *latrophilin cirl* [g14495] were expressed at the anterior border in each segment. The cluster 9 markers, *Awh* [g21739], g15398 (assayed here), *Emx3* [g27623] and *Emx4* [g27624] [46], all show metameric expression from the cheliceral to opisthosomal segments in ring like patterns around the proximal base of each appendage at stage 9 (Fig 5A). Cluster 9 is also marked by *Dfd-A* indicating that it may more specifically correspond to the prosomal leg-bearing segments (Fig 5A). However, this may coincide with the earlier stage 8 expression of *Awh* and g15398, which are predominantly expressed in the prosoma.

The overlap between clusters 1, 7 and 9 may be due to issues regarding the broad distribution of cluster 1 cells that could disrupt marker prediction power. However, given the broad spatial expression of marker genes for clusters 7 and 9, these cells are likely also distributed across the embryo. Given the function of *ham* in the PNS, *Irx* genes in neural development [reviewed in 66] and *Awh* in (motor) neuron cell types, these clusters possibly relate to PNS cells that innervate appendages [67–70].

### New resolution of segment addition and maturation

A key process during stages 7 to 9 is the generation of most of the body segments. The precheliceral, cheliceral and pedipalp segments are instructed through wave-splitting, whereas leg bearing segments of the prosoma are generated by the subdivision of the central region of the germ band [50]. However, the twelve opisthosomal segments are added sequentially from the SAZ that develops from the posterior region of the germ disc and caudal lobe [25]. While candidate gene approaches and screens have provided insights into the regulation of opisthosomal segment generation from the SAZ [21, 25, 26], we explored our single cell data to try to better understand this further during stages 7 to 9.1.

We observed that posterior Hox genes have highest expression in clusters 3 and 10 (Fig 3A). We then examined whether genes known to be expressed in the SAZ were also markers of these clusters. We found that *Wnt8* [g19404], *Wnt11-2* [aug3.g1356], *hh*, and *even-skipped* (*eve*) [g21109], were all markers of cluster 3, while *caudal* (*cad*) was also expressed in cluster 3 (Fig 6A) [22, 25, 26, 71, 72]. We also assayed the spatial expression of markers *AP2* [aug3.g23531] and *g30822* [aug3.g27670], which had been previously reported, and corroborated their expression in the SAZ (Fig 6B) [50]. Furthermore, we analysed the spatial expression of four additional markers of cluster 3 with previously unknown expression, *RNF220* [g27156], *dentin-like*/*DSPP* [g3028], *big-brother* [g8835] and *band4.1* [g22446] (Fig 6B). Like *Wnt8*, *AP2* and g30822 [26, 50], the expression of *big-brother* and *band4.1* was restricted to the posterior SAZ cells (Fig 6B). In contrast, *RNF220* and *DSPP*, like *hh* and *eve* [22, 25], had dynamic expression in the SAZ and in stripes anteriorly in forming segments (Fig 6B). Therefore, we were able to identify new genes from cluster 3 with posteriorly restricted expression that likely maintain the SAZ and regulate segment formation, and genes with cyclical expression associated with the sequential formation of new segments.

**Fig 6:**
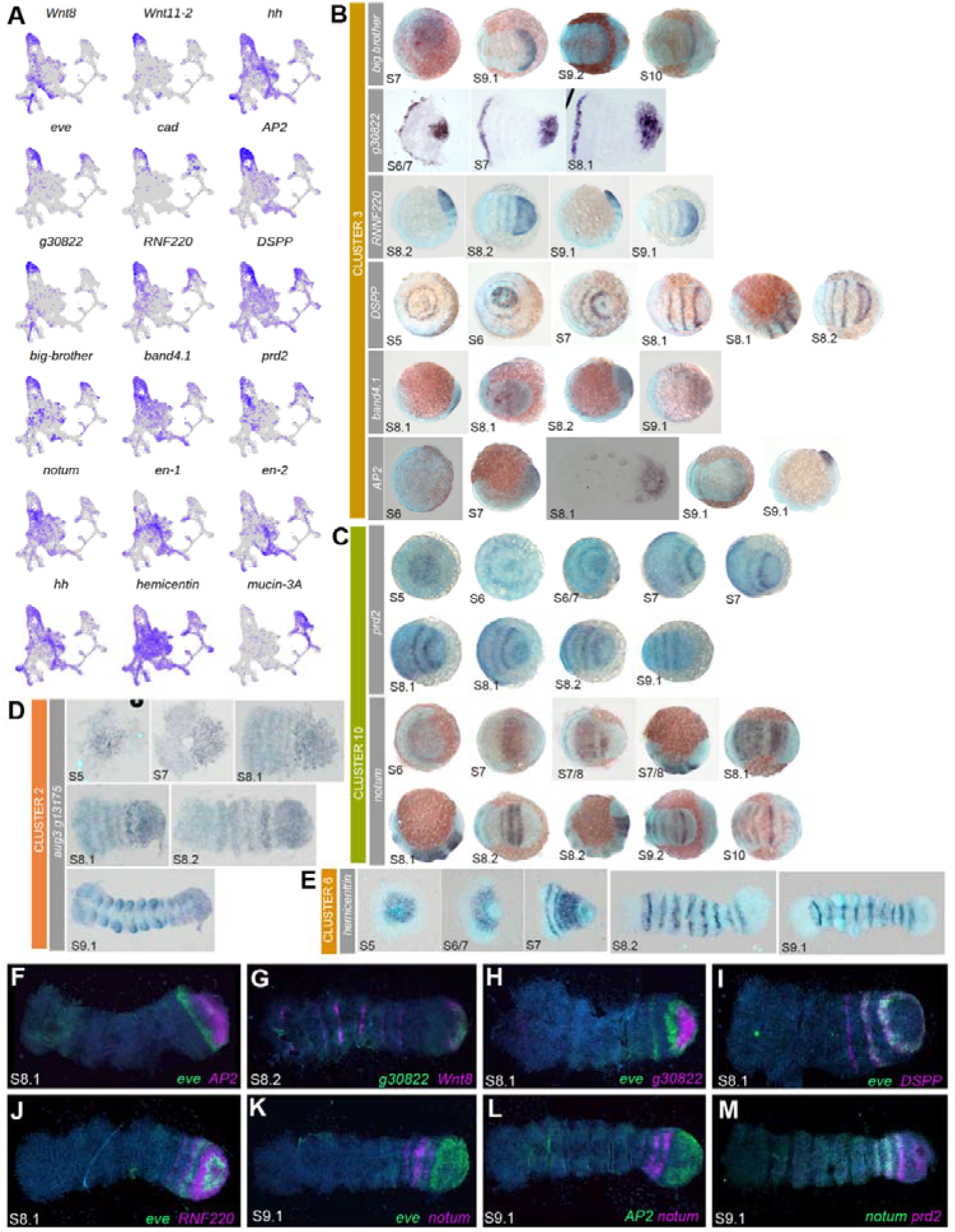
Multiple clusters relating to the SAZ and maturation of posterior segments. (**A**) UMAPs of markers and previously identified genes expressed in the SAZ. (**B** – **E**) Single ISHs of markers for cluster 3 (**B**), for cluster 10 (**C**), for cluster 2 (**D**) and for cluster 6 (**E**). (**F** – **M**) Double FISH, showing; (**F** - **H**) static genes *Wnt8*, *AP2*, and g30822; (**I**) dynamic gene *DSPP* in phase with *eve*, and (**J**) dynamic gene *RNF220* out of phase with *eve*; (**K** – **M**) *notum* and *prd2* expression in relationship each other (**M**) to *eve* (**K**) and to *AP2* (**L**).

We analysed the expression of two cluster 10 markers *paired2* (*prd2*) [g10589], and *notum* [g17362] (Fig 6C) [73]. Note that *gsb*/*prd2* expression up to stage 8 was reported previously [50]. While these two genes were expressed in many regions of the embryo, they had similar expression in the opisthosomal region anterior to the SAZ, like both *Antp* genes at stage 9 (Fig 6C) [14]. Additionally, *prd2* was also expressed in the posterior region of the SAZ (Fig 6C)

To better understand the relationship and phasing of markers from these two clusters we assay the relative expression of marker genes from cluster 3 and 10. Some genes from cluster 3, like *Wnt8*, *AP2* and g30822, had overlapping expression restricted to the posterior of the SAZ, albeit with different anterior borders (Fig 6F – 6H). Interestingly, we observed that some *AP2* and g30822 expressing cells were mutually exclusive from *eve*, which suggests that cluster 3 includes cells that are no longer in the domain of *Wnt8*/*AP2*/g30822 in the posterior SAZ but are in the striped domains more anterior (Fig 6F – 6H). Genes like *eve*, *DSPP* and *RNF220* were all dynamically expressed, however, *DSPP* and *RNF220* exhibited different phasing relative to *eve* (Fig 6I & 6J). This suggests cluster 3 may represent cells across (a) whole segment(s), with different gene expression profiles prefiguring their sub-segmental A-P position (Fig 6I & 6J). The expression of genes from cluster 10 overlapped (e.g. *notum* and *prd2*) but were mostly spatially distinct from cluster 3 genes (Fig 6K – 6M), for example neither *AP2* nor *eve* expression overlapped with *notum*. Therefore, cluster 10 likely represents cells of a transitioning zone where nascent segments have formed anterior to the SAZ and are maturing, which is consistent with *Antp* expression (Fig 6K – 6M) [14].

During posterior segment formation, *engrailed* (*en*) expression is initiated at the parasegmental boundaries and posterior compartments of formed segments anterior to the SAZ [2, 22, 74, 75]. We found both *P. tepidariorum en* paralogs [g24362 and aug3.g15983] were markers of cluster 6, which are also marked by *hh* and another gene not studied so far, *hemicentin* [g1871] (Fig 6E). *hh* and *hemicentin* are also expressed in the posterior compartment of all segments (Fig 6E) [22, 50]. This suggests cluster 6 represents cells in segments later during segmentation than in cluster 3 and 10. However, they only represent a subset of cells in each segment, and likely also relate to both prosomal and opisthosomal cells.

During assessment of markers, we observed that several cluster 3 markers, including *eve*, *DSPP* and *RNF220*, were also cluster 2 markers or at least expressed in cluster 2 (Fig 6A). Cluster 2 was also marked by genes such as *SoxD2* [g19045] known to be expressed in the mesoderm of opisthosomal segments [49], as well as *FGFR1* [13], *integrin-alpha-8* [39] and *ush*, which were markers of dorsal and mesodermal clusters shown previously (Fig 2B). In addition, *twist* was expressed in cluster 2 cells, supporting that at least some of these cells are mesodermal (Fig 2C) [16]. We analysed another cluster 2 marker, *mucin-3A* [aug3.g13175] and found that it was also expressed in the SAZ region and more anteriorly (Fig 6D). This suggests that cluster 2 may represent cells that are becoming internalised posteriorly and contribute to the mesoderm as segments are added [16, 17, 26].

Collectively, our data suggest cell clusters 2, 3, 6 and 10 are associated with posterior segmentation. These clusters relate to the SAZ and other cells more anteriorly that represent differentiating cells in formed segments. Furthermore, the cross over between SAZ and mesodermal markers is consistent with the generation of multiple cell layers during posterior segmentation.

## Discussion

### A combination of ACME and SPLiT-seq for scRNA-seq developmental analysis

This study reports for the first time that a combination of ACME dissociation [29] and SPLiT-seq [40] is a very successful approach to obtain robust single cell data from arthropod embryos. ACME dissociation uses fixation of samples early in the dissociation process allowing us to collect multiple stages and samples that could be stored prior to single cell transcriptomics. This was highly desirable since embryogenesis progresses quickly in many arthropods and capturing distinct stages using lengthy (∼hours) dissociation processes might alter the single-cell expression. Furthermore, unfixed dissociated embryonic cells acquire stress altering their gene expression. Fixing embryos and cells with ACME stopped biological activity, therefore overcoming these hurdles, making the approach highly suited to study embryogenesis. We combined ACME with a variation of SPLiT-seq to obtain tens of thousands of single-cell transcriptomic profiles from multiple embryonic stages. This enabled us to capture new information about the genetic regulation and specification of A-P, D-V and head patterning, mesodermal and endodermal lineages, the CNS and PNS, and posterior segmentation. Therefore, the combined application of ACME and SPLiT-seq is a powerful method to robustly profile development.

### A single-cell atlas of spider development

Our scRNA-seq analysis of the spider *P. tepidariorum* has enabled a new understanding of spider embryogenesis at cellular resolution, including previously uncharacterised genes and unknown aspects of development. We used the integration of three developmental stages to define cell clusters that were represented by cells from all stages (Fig 1). The assessment of markers between each stage and integrated data suggested that this approach did not mask stage specific cell clusters (Sup Fig 5 & 6), implying that between stage 7 to 9.1 no completely distinct cell states emerge. From the integrated data we defined 20 clusters, which represented known as well as novel cell states in multiple aspects of spider embryogenesis. We found multiple previously characterised genes in the clusters, which allowed us to establish their identities. Most cell clusters were also marked by genes that had not been previously studied and therefore offer new insights into these cells and the regulation of developmental processes. Collectively, our dataset constitutes the first cell state atlas through spider embryonic stages 7 to 9.1.

We detected fundamental aspects of developmental regulation relating to Hox patterning of the A-P axis (Fig 3), the formation of the D-V axis (Fig 2 & 5) and germ layer cell types (Fig 2). These aspects reveal that the genetic components that regulate these core geometries also regionalise/cluster cells in our scRNA-seq data. The Hox genes were clearly segregated into three Hox positive groups, relating to pedipalpal, leg bearing and opisthosoma segments (Fig 3). Yet within each of these regions, ohnologs appeared to show differential divergences that might relate to sub- and/or neo-functionalisation, indicating a potential for scRNA-seq to study general trends of ohnolog evolution after WGD in a large scale (Fig 3 & Sup Fig 7). The mesodermal/endodermal clusters carried a signature of increased G1 proportion, which suggests less proliferation compared to ectodermal clusters (Fig 2). This potentially reflects the early determination of these cell types in spider development as shown in single cell data from earlier in embryogenesis [39] and that they are transcriptional very distinct at stages 7 to 9.1.

### Extra-embryonic cells contribute not only to the gut but also hemocytes

The germ disc and germ band of spiders has been largely considered to be exclusive from the extra embryonic region and yolk. However, gut genes *serpent* and *hnf-4* [48], and the endodermal marker *GPCPD* [12], are expressed in the so called extra-embryonic region, suggesting these cells contribute to the spider endoderm. These genes were present in cluster 20, along with previously unknown markers that were also expressed in the extra-embryonic region (Fig 2). By cell cycle scoring of clusters, we revealed that these cluster 20 cells are likely dividing less than many germ band related cell clusters.

While these endodermal cells related to cluster 20 have been previously reported, the expression of markers in the extra-embryonic region from cluster 19 suggests a need for reinterpretation of this tissue beyond only contributing to the gut. Prior to this study, the mesodermal heart gene *tinman* had been investigated in the spider *Cupiennius salei* [76], however, nothing was known about the genetic specification of peripheral circulatory cells in spiders. We found that cluster 19 expressed *Mef2.1*, which was later expressed in the heart (Fig 2), as well as *hemocyanins.* This strongly suggests that cluster 19 may relate to heart, hemolymph and hemocyte cells. These cells initially surround the dorsal precheliceral region, but subsequently migrate into the extra embryonic region. Interestingly, in *D. melanogaster*, embryonic hemocytes also originate from head mesoderm suggesting a perhaps conserved origin between chelicerates and insects [57–60]. In *D. melanogaster* and *Tribolium castaneum*, the extra-embryonic tissue is involved in immune response, and our identification of hematopoiesis markers suggest for the first time that cells in the extra-embryonic region of spider embryos could also contribute to this function [77, 78]. Thus, our data offer the opportunity to investigate blood/immune system evolution and function.

### Strong signal of D-V patterning in *P. tepidariorum* scRNA-seq

Like other arthropods, *P. tepidariorum* D-V axis formation and patterning is regulated by ventral *sog* and dorsal *dpp* signals [8, 9, 79]. In addition, genes like *Ets4*, *hh* and *fgf8* are also known to control cumulus migration, a process which is crucial for spider D-V axis formation [12, 13, 80]. However, there is still much to understand about the evolution of spider D-V patterning.

We observed a strong ventral signature of *sog* expression across cell clusters associated with the ventral germ band proper (Fig 5). As expected, *sog* and other cluster 8 markers, like *Nkx6.2*, were expressed along the ventral midline, as in other animals [65]. Clusters 7 and 9 were also marked by *sog* and other markers of these clusters exhibited complex expression likely associated with the PNS, given the function of genes like *hamlet*, *awh*, *Emx*, *Irx* and *netrin* [66–70]. Therefore, our single cell data revealed that alongside fundamental aspects of ventral specification, stages 7 to 9.1 also include patterning of the ventral PNS.

With respect to dorsal specification and patterning, genes expressed around the dorsal rim of the germ-band marked cluster 5 along with *dpp* (Fig 2). Furthermore, there were differences between cell cycle scoring between these dorsal and ventral clusters, implying dorsal cells were not proliferating as much as ventral cells (Fig 2). Curiously, we observed *noggin-D* expression in the cumulus, dorsal field, and dorsal region of the germ band (Fig 2). The *Xenopus leavis noggin* homolog is expressed dorsally with *chordin*/*sog* [81] and ectopic expression can ventralise *D. melanogaster* embryos, thus revealing its conserved function as a BMP signalling inhibitor [81]. It is therefore surprising that *P. tepidariorum* exhibits dorsal expression of *noggin-D* where *dpp* signalling is active and necessary for dorsalisation [8]. This suggests that while *sog* and *dpp* play conserved roles in D-V specification in *P. tepidariorum,* other genes like *noggin-D* have diverged in function, indicating potential developmental system drift of D-V regulation in *P. tepidariorum*.

### New insights into precheliceral region patterning

During prosomal development, *otd-1* helps specify the precheliceral region [45], where structures/organs such as the brain, eyes and stomodeum develop. Markers of clusters 13, 14 and 16 showed specific spatial expression suggestive of demarcation of different pre-cheliceral regions (Fig 4). For example, expression of *six3* paralogs in cluster 14 remained at the anterior rim and suggests conserved roles for these genes in forebrain development [64]. Furthermore, in several insects and the centipede *Strigamia maritima*, the combination of *six3* and *vsx* denotes the region of the anterior medial region that forms the *pars intercerebralis* [82–85]. Since *P. tepidariorum six3* paralogs and *vsx* are expressed in cluster 14 and anterior of the precheliceral region, there is a possibility that *P. tepidariorum* shares a conserved regulatory control of *pars intercerebralis*. However, while neurogenic clusters were identified we were unable to definitively relate clusters to the non-neurogenic ectoderm that also forms at the anterior and lateral rim of the precheliceral region [18, 19]. Future, identification of these cells could give further insights into differentiation of the non-neurogenic ectoderm and eye development.

Expression of clusters 13 and 16 markers were observed initially at the anterior but subsequently shifted posteriorly to different extents and separated along the D-V axis (Fig 4). Markers of these clusters, *lim1a*, *Pax6.1* and *Pax6.2*, are known for their essential roles in head and neural development in other arthropods and other animals [86–88]. Overall, our scRNA-seq reveals three cell clusters present from stage 7 that likely prefigure different regions of the head and brain (Fig 4).

### Regionalisation of posterior segmentation based on genetic signatures of segment formation and maturation

Although previous studies identified key genes and their interactions regulating posterior segment addition in spiders and other arthropods [17, 25, 26, 89–95], we still have a poor understanding of how SAZs work. There is evidence that SAZs are likely genetically sub-structured, representing different regions and/or states that cells must progress through to form new segments [25, 96]. Our approach has identified more robust genetic signatures of SAZ sub-structure and revealed new genes involved in segmentation. This allows us to propose an extended model for the structure of the SAZ and segment addition.

Our data suggest that opisthosomal segmentation can be divided into four regions of segment formation (Fig 6): region one relates to the most posterior cells in the SAZ, marked by genes like *Wnt8, AP2* and g30822 that have static expression [50]. These cells and several of the markers including *Wnt8*, can already be observed in stage 5 single cell data consistent with the SAZ forming from the caudal lobe during stages 5 and 6 [39]. As previously suggested, this posterior region of the SAZ probably represents a pool of undifferentiated cells that continuously contributes to new segments, perhaps analogous to the caudal region of the vertebrate presomitic mesoderm [26, 97].

Anteriorly in the SAZ, region two is marked by phased expression of pair-rule gene orthologs, like *eve* and *runt* [25], and newly identified genes including *RNF220* and *DSPP* (Fig 6). These genes don’t appear to mark cell clusters at stage 5, which is consistent with these cell states only developing as the SAZ starts to generate segments during stages 6 and 7 [39]. These phased domains appear to broadly relate to A–P regions within forming segments (Fig 6) [22, 25, 43, 50]. Interestingly, *RNF220* enhances canonical Wnt signalling in other animals, and therefore might modulate the *Wnt8* activity in the SAZ of *P. tepidariorum* [98]. Region two therefore likely represents forming segments anteriorly from the SAZ, but still lacking expression of the segment polarity genes like *en* (Fig 6) [2].

Region three is marked by expression of *notum,* which is also expressed in a similar pattern in *C. salei* [99]. *notum* is expressed in the posterior region of segments in this region (Fig 6). *notum* is a Wnt suppressor and so it may act on Wnt activity to facilitate segment maturation [26, 73]. Due to this transition from the SAZ to a more differentiated region we describe this region as the segment maturation zone (SMZ) (Fig 6). Additionally, the expression of *sog* in cluster 10 also suggests that segmental patterning along the D-V axis is regulated anterior to the SAZ.

Formed segments express segment polarity genes like *en*, which marks the fourth region (Fig 6) [2]. Indeed our data identified a clear genetic signature of cells in the posterior compartment of segments [75, 100, 101]. This includes expression of *hemicentin,* which is also expressed in developing somites of zebrafish [102], and knockdowns exhibit detachment phenotypes between cell types. Therefore, maturation of spider posterior segments may also involve Hemicentin mediated inter-cellular interactions between cells of multiple clusters defined in this study.

The SAZ cluster 3 is marked by several genes also expressed in cluster 2 (Fig 6). Cluster 2 corresponds to mesodermal cells, given the expression of *SoxD2*, *FGFR1* and *integrin-alpha-8*, which are also mesodermally expressed in other animals [49, 103–105]. Both *integrin-alpha-8* and *SoxD2* are expressed in the SAZ and anterior to it, but also later are expressed in mesodermal metameric blocks in the opisthosoma and prosoma [49]. This is reminiscent of the *SoxD2* vertebrate homolog, *Sox5*, which is expressed in the presomitic mesoderm and later within each formed somite [103]. Furthermore, disruption of the SAZ by RNAi against *Wnt8* or *Dl* results in ectopic expression of the mesoderm gene *twist* in the SAZ [16, 17, 26]. This strongly suggests that there is dynamic specification and sorting of ectodermal and mesodermal cells via cell movement in the SAZ [104, 105]. Another marker of cluster 2, *Ubx-A*, has been shown to suppress *twist* in somatic myogenesis, which raises the possibility that this Hox gene may regulate ectoderm-mesoderm dynamics in the spider SAZ [106].

## Conclusion

Our scRNA-seq cell atlas of spider development corroborates previous findings and also provides novel insights into several important processes during spider embryogenesis. Given that we only examined three embryonic stages and one earlier stage has been examined recently (ref), there is still a wealth of information awaiting capture from scRNA-seq at other time points in spider development. Future work to compare spider cell atlases to those of other chelicerates and other arthropods will also provide new insights into the evolution of cell differentiation and fate as well as the regulation of embryogenesis more broadly.

## Materials and Methods

### Dissociation of *P. tepidariorum* embryos for single-cell sequencing

*P. tepidariorum* embryos were collected from an inbred culture, staged [6] and dissociations were performed as previously described [29]. Stage 7 (51-55 hours after egglaying), 8.1 (56-55 hours after egglaying) or 9.1 (76-80 hours after egglaying) embryos were selected and weighed without silk to determine the sample size. Embryos were dechorionated with bleach (sodium hypochlorite, 5% active chlorine, Arcos) and tap water (1:1). Embryos were washed several times with ultrapure water (UltraPure™ DNase/RNase-Free Distilled Water, Invitrogen) to remove bleach traces. Unfertilized embryos were removed, and embryos were immersed in 10 ml ACME solution (3:1:2:14 of methanol, glacial acetic acid, glycerol, and ultrapure water). To break open the vitelline membranes and allow complete dissociation, embryos were treated with a few pulses of polytron homogenisation. Embryos in ACME solution were incubated for 1 hour at room temperature (RT) on a rocking platform (Stuart SSL4) at 70 oscillations per minute. The cell suspension was filtered through a 50 μm filter (Sysmex/Partec CellTrics, Wolflabs) to separate remaining cell clumps and debris (vitelline membranes). Dissociated cells were pelleted at 1500 rpm for 5 minutes at 4°C. The supernatant was discarded, and the cell pellet washed with 7 ml PBS/1% Bovine Serum Albumin (BSA) (BSA Microbiological Grade Powder, Fisher BioReagents). The cells were pelleted at 1500 rpm for 5 minutes at 4°C, the supernatant was discarded, and the cell pellet resuspended in 1 ml 1x PBS-1% BSA and DMSO (9:1) and stored at -20°C.

### Flow cytometry and cell dilution

ACME-dissociated cells from stage 7, 8 and 9 embryos were thawed on ice. Thawed cells were centrifuged twice at 1500 rpm for 5 minutes at 4°C, to remove the DMSO, and resuspended in 400 μl of fresh 1x PBS-1% BSA. Samples were then filtered through a 50 μm filter (Sysmex/Partec CellTrics, Wolflabs) and collected into new 1.5 ml Eppendorf tubes, on ice. 50 μl of filtered cells were added to 100 μl of 1x PBS-1% BSA. The remaining undiluted samples were kept on ice, in the fridge, for the rest of the analysis.

Dilutions were stained with 0.4 μl of DRAQ5 (5 mM stock solution, Bioscience) and 0.8 μl of Concanavalin-A conjugated with AlexaFluor 488 (1 mg/ml stock solution, Invitrogen), and incubated in the dark for 25 min at room temperature. DRAQ5 was used as nuclear dye, while Concanavalin-A (Con-A) was used as cytoplasmic dye. We visualized and counted cells using a CytoFlex S Flow Cytometer (Beckman Coulter). For each stained dilution, we made three measurements of 10 μl and registered the average number of total events (total ungated population). From this we calculated the number of total events per μl in our undiluted samples.

When multiple samples were available, we selected those with the highest percentage of singlets (DRAQ5-positive & Concanavalin-A-positive single-cells). To obtain this percentage of singlets, we used the following gating strategy: FSC-H vs FSC-A, where we selected only well-correlated events (first filter to remove aggregates); Con-A vs FSC-A, where we selected Con-A positive events (events with cytoplasm); DRAQ5 vs FSC-A, where we selected DRAQ5 positive events (events with nucleus); DRAQ5-A vs DRAQ5-H, where we selected only well-correlated events (second filter to remove aggregates); and DRAQ5 vs Con-A, where we obtained the final number of singlets.

To prepare the cells for the SPLiT-seq protocol, we unstained the undiluted/unstained samples. Samples were diluted in fresh 0.5x PBS buffer to a final concentration of 625 events/μl and kept on ice.

### Re-annotation of *P. tepidariorum* genome for mapping SPLiT-seq data

SPLiT-seq has a bias towards capturing the 3’ region of transcripts. To ensure capture of the signal from the SPLiT-seq data we re-annotated the genome (GCA000365465.2) of *P. tepidariorum*. Bulk RNA-seq data [107] was combined with multiple paired end libraries from a range of other embryonic stages. All data were quality trimmed with Trimmomatic v0.39 [108] and then mapped to the genome using Star v2.7.9a [109] using the 2-pass method for better detection of splice junctions. Alignment information was used as evidence for Braker v2 [110] annotation, with ten rounds of optimisation, UTR training and considering CRF models. This annotation was combined with the previous genome annotation [14] by first merging the gene coordinates with bamtools merge. Gene models with a new annotation were replaced. Those with multiple new annotations to one previous annotation were rejected and the previous annotation was retained. New annotations that compounded multiple previous annotations were retained. This final annotation contained 33,413 gene models compared to the 27,950 in the Schwager et al. (2017) annotation. The majority (18,544) of these genes show a 1:1 relationship between annotations, as well as additional annotations captured by the new version, and the fusion of split genes by merging annotations. Old annotations are given as aug3.g* whereas new annotations are given as g*. The reannotation GTF and amino acid fasta files have been uploaded to figshare (https://doi.org/10.6084/m9.figshare.c.6032888.v2). Gene orthologues were estimated by alignment scoring using Diamond tblastx [111, 112] to the NCBI nr database.

### Mitochondrial genome assembly of *P. tepidariorum*

Mitochondrial expression in single-cell sequence data can be indicative of cell stress and therefore a useful metric to measure. We assembled a partial, but near complete, version of the mitochondrial genome to be included in the mapping steps. DNA-seq data (SRR891587) from the genome assembly was trimmed with Trimmomatic v0.39 [108] and assembled with Spades v3.13.1 [113] using kmer sizes 21, 33, 55, 77 with the *P. tepidariorum* CO1 sequence (DQ029215.1) as a trusted contig. The spades contig matching the CO1 sequence was extracted and then extended with NOVOPlasty v4.2 [114] to achieve a final assembly of length 14,427 bp, though it was not circularisable. MiToS v2 [115] identified all expected features. The sequence was added to the genome assembly and a feature spanning the full length was added to the GTF gene coordinates file for mapping. The mitochondrial genome assembly has been uploaded to figshare (https://doi.org/10.6084/m9.figshare.c.6032888.v2).

### SPLiT-seq, filtering, pre-processing, and clustering analysis

The SPLiT-seq protocol was performed as previously described with some modifications (Sup Text) [29]. Libraries were sequenced with 150 bp paired-end Illumina NovaSeq 6000 S4 flow cell, provided commercially by Novogene. Raw reads have been uploaded to the ENA with BioProject PRJEB53350.

Total sequencing output was 103.9 + 40.7 Gb, constituting a total of 963,482,454 raw reads, with >99.98% clean reads and a Q20 >93.94%. All data and samples passed FastQC inspection. Adapters and low-quality bases were trimmed with Cutadapt v1.18 [116] and properly paired reads were combined with Picard FastqToSam v2.20.5. All sequence runs were combined with Picard MergeSamFiles to attain paired reads for downstream expression analysis. To generate reference files, first Picard CreateSequenceDictionary was used to generate a dictionary from the genome plus mitochondrial sequence and re-annotations. Then converted to a RefFlat, a reduced GTF and intervals with DropSeq v2.4.0 [117] tools ConvertToRefFlat, ReduceGtf and CreateIntervalsFiles, respectively. For mapping the data, a Star v2.7.9a [109] genome index was generated with sjdbOverhang 99. These reference files and genome index were used as inputs for the Split-seq_pipeline.sh [117]. An expression matrix was generated with dropseq DigitalExpression, including reads mapping to introns, with a barcode edit distance of one, and outputting cells that had at least 100 genes. Cells from each stage were extracted from this matrix using the 16 sequences from cell barcode one.

Each library per stage was first processed for doublet removal. The expression matrix for each stage was loaded into Seurat v4 [52] and subset to contain cells where genes are expressed in at least 20 cells; that have genes numbers between 400 and 1800; UMI counts between minimums of 650 for stage 7, 700 for stage 8.1 and 500 for stage 9.1 and maximum of 4500; and no more than 1% mitochondrial expression. This initial dataset contained cells for stages 7 (1967 and 3111), 8.1 (2058 and 3029) and 9.1 (3866 and 5459), for libraries one and two respectively. Each sample was normalised with SCTransform with the glmGamPoi method and variable features threshold of 1.3 and regressing the mitochondrial expression, UMI counts and gene counts. 50 PCs and neighbours were computed using k. param 100, and clusters were identified at a resolution of 1 for stage 7 and 8.1 and 1.2 for stage 9.1. Using doubletFinder v3 [118], 5% doublets we removed, identifying an appropriate pK with an initial parameter sweep, and retained singlets were extracted.

Doublet filtered stage specific matrices were then processed for integration in Seurat, normalising with SCTransform using the “glmGamPoi” method and a variable feature threshold of 1.3 and regressing the mitochondrial expression, UMI counts and gene counts. All samples were integrated using the reciprocal PCA method (50 PCs) and 40 anchors. Fifty PCs were computed for the integrated data and used for UMAPs with the “umap-learn” method and 100 n.neighbors, 0.3 min.dist, 42 seed and the “correlation” metric. Nearest neighbours were determined using 100 k.param and 50 n.trees. The Leiden algorithm was used for clustering with a resolution of 1.2. The clustering resolutions were guided by ChooseR [119] and clustree [120] analysis. FindAllMarkers was used to extract markers for each cluster using the Wilcoxon method and including genes that were expressed in at least 25% of the cells in their respective cluster and a return threshold of 1e-5. Marker genes were annotated initially with the NCBI nr database using Diamond v2.0.8.146 [111, 112] and refined for existing genes already characterised in *P. tepidariorum*. Hierarchical grouping of clusters was performed using BuildClusterTree from Seurat and ggtree [121] to visualise. Seurat objects available upon request.

### Clustering and marker comparisons

Clustering from different Seurat processing iterations were compared using the adjustRandIndex from the R package mclust [53]. Raw count matrices were processed similarly with a variable gene threshold, ranging from 1.2 to 1.7, and the integration k.anchor rnaging from 5 to 45. Cluster IDs were extracted and for an all versus all comparison of clustering similarity scored between 0 and 1 and were visualised with pheatmap in R. UMAP coordinates were extracted and plotted with ggplots in R.

To compare marker lists of clusters from different runs, FindAllMarker was used with 25 percent expressed in cluster and 1e-5 *p*-value threshold parameters and compared using a hypergeometric distribution test. The total number of genes in the spider single cell expression matrix was used as the pool from which genes could be selected. The markers and overlap were computed from each cluster, and these values were used in a sum(dhyper()) R function with a Bonferroni adjusted *p*-value. Code for the hypergeometric distribution test can be found at https://github.com/djleite/Hypergeo_SingleCell_Markers.

### Cell Cycle Scoring

The cell cycle scores of *P. tepidariorum* were estimated in Seurat. To identify cell cycle genes the G2/M and S phase *Drosophila melanogaster* gene IDs were obtained from (https://github.com/hbc/tinyatlas/blob/master/cell_cycle/Drosophila_melanogaster.csv) and extracted from the release v6.49 of the *D. melanogaster* proteins from FlyBase. Spider orthologs were identified using a default Diamond blastp [111, 112] search, selecting two best hits. Spider cell cycle gene IDs were used with Seurat CellCycleScoring to identify cell cycle phasing. Fasta sequence of *P. tepidariorum* cell cycle genes are provided in Sup File 2.

### Gene cloning and expression analysis

For gene expression characterisation in *P. tepidariorum* embryos we performed colorimetric *in situ* hybridisation (ISH) [122], fastred [20] and double fluorescent in situ hybridisation (dFISH) [49] as previously described with minor modifications (Sup Text).

## Ethics approval and consent to participate

Not applicable

## Consent for publication

Not applicable

## Availability of data and materials

Raw reads for the scRNA-seq have been uploaded to the ENA with BioProject PRJEB53350. The re-annotation of the genome, the mitochondrial genome assembly of *P. tepidariorum* and the scRNA-seq expression matrices have been uploaded to figshare (https://doi.org/10.6084/m9.figshare.c.6032888.v2).

## Authors’ contributions

APM, AS, DJL and JS conceptualised the study. AS and HGC performed the single-cell sequencing. DJL and NJK performed the bioinformatic analysis. DJL developed software for analysis. GB, AH, MP, NT, LBG, RW, NS, AGN and VSPK performed in situ expression analysis. DJL, APM and JS wrote the main manuscript. All authors reviewed the manuscript.

## Acknowledgements

We thank Lauren Sumner-Rooney and Fritz Vollrath for discussions. We thank Helen Ferry and Liam Hardy at the Experimental Medicine Division Flow Cytometry Facility at the Nuffield Department of Clinical Medicine (University of Oxford), Michal Maj, and Robert Hedley at the Flow Cytometry Facility at the Dunn School of Pathology (University of Oxford) for their help and advice with flow cytometry. We thank Vincent Mason for technical help. We thank the Center for Advanced Light Microscopy at the LMU Munich and Gregor Bucher for providing the microscope set-up for imaging whole mount in situ hybridisation.

## References

1. Carroll SB, Grenier JK, Weatherbee SD. From DNA to diversity: Molecular genetics and the evolution of animal design. Malden, Mass: Blackwell Science; 2001.

2. Schwager EE, Schöenauer A, Leite DJ, Sharma PP, McGregor AP. Chelicerata. In: Wanninger A, editor. Evolutionary Developmental Biology of Invertebrates 3: Ecdysozoa I: Non-Tetraconata.: Springer-Verlag; 2015.

3. Oda H, Akiyama-Oda Y. The common house spider Parasteatoda tepidariorum. Evodevo. 2020;11:6. Epub 20200320.

4. McGregor AP, Hilbrant M, Pechmann M, Schwager EE, Prpic NM, Damen WG. Cupiennius salei and Achaearanea tepidariorum: Spider models for investigating evolution and development. Bioessays. 2008;30(5):487–98.

5. Hilbrant M, Damen WG, McGregor AP. Evolutionary crossroads in developmental biology: the spider Parasteatoda tepidariorum. Development. 2012;139(15):2655–62.

6. Mittmann B, Wolff C. Embryonic development and staging of the cobweb spider Parasteatoda tepidariorum C. L. Koch, 1841 (syn.: Achaearanea tepidariorum; Araneomorphae; Theridiidae). Dev Genes Evol. 2012;222(4):189–216. Epub 20120509.

7. Pechmann M. Formation of the germ-disc in spider embryos by a condensation-like mechanism. Front Zool. 2016;13:35. Epub 20160811.

8. Akiyama-Oda Y, Oda H. Axis specification in the spider embryo: dpp is required for radial-to-axial symmetry transformation and sog for ventral patterning. Development. 2006;133(12):2347–57.

9. Akiyama-Oda Y, Oda H. Early patterning of the spider embryo: a cluster of mesenchymal cells at the cumulus produces Dpp signals received by germ disc epithelial cells. Development. 2003;130(9):1735–47.

10. Hemmi N, Akiyama-Oda Y, Fujimoto K, Oda H. A quantitative study of the diversity of stripe-forming processes in an arthropod cell-based field undergoing axis formation and growth. Dev Biol. 2018;437(2):84–104. Epub 20180316.

11. Oda H, Iwasaki-Yokozawa S, Usui T, Akiyama-Oda Y. Experimental duplication of bilaterian body axes in spider embryos: Holm’s organizer and self-regulation of embryonic fields. Dev Genes Evol. 2020;230(2):49–63. Epub 20190410.

12. Akiyama-Oda Y, Oda H. Cell migration that orients the dorsoventral axis is coordinated with anteroposterior patterning mediated by Hedgehog signaling in the early spider embryo. Development. 2010;137(8):1263–73.

13. Wang R, Leite DJ, Karadas L, Schiffer PH, Pechmann M. FGF signalling is involved in cumulus migration in the common house spider Parasteatoda tepidariorum. Dev Biol. 2022;494:35–45. Epub 20221205.

14. Schwager EE, Sharma PP, Clarke T, Leite DJ, Wierschin T, Pechmann M, et al. The house spider genome reveals an ancient whole-genome duplication during arachnid evolution. BMC Biol. 2017;15(1):62. Epub 20170731.

15. Pechmann M, Schwager EE, Turetzek N, Prpic NM. Regressive evolution of the arthropod tritocerebral segment linked to functional divergence of the Hox gene labial. Proc Biol Sci. 2015;282(1814).

16. Yamazaki K, Akiyama-Oda Y, Oda H. Expression patterns of a twist-related gene in embryos of the spider Achaearanea tepidariorum reveal divergent aspects of mesoderm development in the fly and spider. Zoolog Sci. 2005;22(2):177–85.

17. Oda H, Nishimura O, Hirao Y, Tarui H, Agata K, Akiyama-Oda Y. Progressive activation of Delta-Notch signaling from around the blastopore is required to set up a functional caudal lobe in the spider Achaearanea tepidariorum. Development. 2007;134(12):2195–205. Epub 20070516.

18. Schomburg C, Turetzek N, Schacht MI, Schneider J, Kirfel P, Prpic NM, et al. Molecular characterization and embryonic origin of the eyes in the common house spider Parasteatoda tepidariorum. Evodevo. 2015;6:15. Epub 20150428.

19. Baudouin-Gonzalez L, Harper A, McGregor AP, Sumner-Rooney L. Regulation of Eye Determination and Regionalization in the Spider Parasteatoda tepidariorum. Cells. 2022;11(4). Epub 20220211.

20. Janeschik M, Schacht MI, Platten F, Turetzek N. It takes Two: Discovery of Spider Pax2 Duplicates Indicates Prominent Role in Chelicerate Central Nervous System, Eye, as Well as External Sense Organ Precursor Formation and Diversification After Neo- and Subfunctionalization Frontiers in Ecology and Evolution. 2022;10.

21. Paese CLB, Schoenauer A, Leite DJ, Russell S, McGregor AP. A SoxB gene acts as an anterior gap gene and regulates posterior segment addition in a spider. Elife. 2018;7. Epub 20180821.

22. Kanayama M, Akiyama-Oda Y, Nishimura O, Tarui H, Agata K, Oda H. Travelling and splitting of a wave of hedgehog expression involved in spider-head segmentation. Nat Commun. 2011;2:500. Epub 20111011.

23. Oda H, Akiyama-Oda Y. Dataset on gene expressions affected by simultaneous knockdown of Hedgehog and Dpp signaling components in embryos of the spider Parasteatoda tepidariorum. Data Brief. 2020;28:105088. Epub 20200103.

24. Tanay A, Sebe-Pedros A. Evolutionary cell type mapping with single-cell genomics. Trends Genet. 2021;37(10):919–32. Epub 20210518.

25. Schonauer A, Paese CL, Hilbrant M, Leite DJ, Schwager EE, Feitosa NM, et al. The Wnt and Delta-Notch signalling pathways interact to direct pair-rule gene expression via caudal during segment addition in the spider Parasteatoda tepidariorum. Development. 2016;143(13):2455–63. Epub 20160610.

26. McGregor AP, Pechmann M, Schwager EE, Feitosa NM, Kruck S, Aranda M, et al. Wnt8 is required for growth-zone establishment and development of opisthosomal segments in a spider. Curr Biol. 2008;18(20):1619–23.

27. Stuart T, Satija R. Integrative single-cell analysis. Nat Rev Genet. 2019;20(5):257–72.

28. Wang J, Sun H, Jiang M, Li J, Zhang P, Chen H, et al. Tracing cell-type evolution by cross-species comparison of cell atlases. Cell Rep. 2021;34(9):108803.

29. Garcia-Castro H, Kenny NJ, Iglesias M, Alvarez-Campos P, Mason V, Elek A, et al. ACME dissociation: a versatile cell fixation-dissociation method for single-cell transcriptomics. Genome Biol. 2021;22(1):89. Epub 20210408.

30. Plass M, Solana J, Wolf FA, Ayoub S, Misios A, Glazar P, et al. Cell type atlas and lineage tree of a whole complex animal by single-cell transcriptomics. Science. 2018;360(6391). Epub 20180419.

31. Briggs JA, Weinreb C, Wagner DE, Megason S, Peshkin L, Kirschner MW, et al. The dynamics of gene expression in vertebrate embryogenesis at single-cell resolution. Science. 2018;360(6392). Epub 20180426.

32. Cao C, Lemaire LA, Wang W, Yoon PH, Choi YA, Parsons LR, et al. Comprehensive single-cell transcriptome lineages of a proto-vertebrate. Nature. 2019;571(7765):349–54. Epub 20190710.

33. Cao J, Packer JS, Ramani V, Cusanovich DA, Huynh C, Daza R, et al. Comprehensive single-cell transcriptional profiling of a multicellular organism. Science. 2017;357(6352):661–7.

34. Cao J, Spielmann M, Qiu X, Huang X, Ibrahim DM, Hill AJ, et al. The single-cell transcriptional landscape of mammalian organogenesis. Nature. 2019;566(7745):496–502. Epub 20190220.

35. Farrell JA, Wang Y, Riesenfeld SJ, Shekhar K, Regev A, Schier AF. Single-cell reconstruction of developmental trajectories during zebrafish embryogenesis. Science. 2018;360(6392). Epub 20180426.

36. Karaiskos N, Wahle P, Alles J, Boltengagen A, Ayoub S, Kipar C, et al. The Drosophila embryo at single-cell transcriptome resolution. Science. 2017;358(6360):194–9. Epub 20170831.

37. Packer JS, Zhu Q, Huynh C, Sivaramakrishnan P, Preston E, Dueck H, et al. A lineage-resolved molecular atlas of C. elegans embryogenesis at single-cell resolution. Science. 2019;365(6459). Epub 20190905.

38. Wagner DE, Weinreb C, Collins ZM, Briggs JA, Megason SG, Klein AM. Single-cell mapping of gene expression landscapes and lineage in the zebrafish embryo. Science. 2018;360(6392):981–7. Epub 20180426.

39. Akiyama-Oda Y, Akaiwa T, Oda H. Reconstruction of the Global Polarity of an Early Spider Embryo by Single-Cell and Single-Nucleus Transcriptome Analysis. Front Cell Dev Biol. 2022;10:933220. Epub 20220722.

40. Rosenberg AB, Roco CM, Muscat RA, Kuchina A, Sample P, Yao Z, et al. Single-cell profiling of the developing mouse brain and spinal cord with split-pool barcoding. Science. 2018;360(6385):176–82. Epub 20180315.

41. Turetzek N, Pechmann M, Schomburg C, Schneider J, Prpic NM. Neofunctionalization of a Duplicate dachshund Gene Underlies the Evolution of a Novel Leg Segment in Arachnids. Mol Biol Evol. 2016;33(1):109–21. Epub 20151006.

42. Turetzek N, Khadjeh S, Schomburg C, Prpic NM. Rapid diversification of homothorax expression patterns after gene duplication in spiders. BMC Evol Biol. 2017;17(1):168. Epub 20170714.

43. Schwager EE, Pechmann M, Feitosa NM, McGregor AP, Damen WG. hunchback functions as a segmentation gene in the spider Achaearanea tepidariorum. Curr Biol. 2009;19(16):1333–40. Epub 20090723.

44. Pechmann M, Khadjeh S, Turetzek N, McGregor AP, Damen WG, Prpic NM. Novel function of Distal-less as a gap gene during spider segmentation. PLoS Genet. 2011;7(10):e1002342. Epub 20111020.

45. Pechmann M, McGregor AP, Schwager EE, Feitosa NM, Damen WG. Dynamic gene expression is required for anterior regionalization in a spider. Proc Natl Acad Sci U S A. 2009;106(5):1468–72. Epub 20090115.

46. Leite DJ, Baudouin-Gonzalez L, Iwasaki-Yokozawa S, Lozano-Fernandez J, Turetzek N, Akiyama-Oda Y, et al. Homeobox Gene Duplication and Divergence in Arachnids. Mol Biol Evol. 2018;35(9):2240–53.

47. Kanayama M, Akiyama-Oda Y, Oda H. Early embryonic development in the spider Achaearanea tepidariorum: Microinjection verifies that cellularization is complete before the blastoderm stage. Arthropod Struct Dev. 2010;39(6):436–45.

48. Feitosa NM, Pechmann M, Schwager EE, Tobias-Santos V, McGregor AP, Damen WGM, et al. Molecular control of gut formation in the spider Parasteatoda tepidariorum. Genesis. 2017;55(5). Epub 20170422.

49. Baudouin-Gonzalez L, Schoenauer A, Harper A, Blakeley G, Seiter M, Arif S, et al. The Evolution of Sox Gene Repertoires and Regulation of Segmentation in Arachnids. Mol Biol Evol. 2021;38(8):3153–69.

50. Akiyama-Oda Y, Oda H. Hedgehog signaling controls segmentation dynamics and diversity via msx1 in a spider embryo. Sci Adv. 2020;6(37). Epub 20200909.

51. Korsunsky I, Millard N, Fan J, Slowikowski K, Zhang F, Wei K, et al. Fast, sensitive and accurate integration of single-cell data with Harmony. Nat Methods. 2019;16(12):1289–96. Epub 20191118.

52. Hao Y, Hao S, Andersen-Nissen E, Mauck WM, 3rd, Zheng S, Butler A, et al. Integrated analysis of multimodal single-cell data. Cell. 2021;184(13):3573–87 e29. Epub 20210531.

53. Scrucca L, Fop M, Murphy TB, Raftery AE. mclust 5: Clustering, Classification and Density Estimation Using Gaussian Finite Mixture Models. R J. 2016;8(1):289–317.

54. Iwasaki-Yokozawa S, Nanjo R, Akiyama-Oda Y, Oda H. Lineage-specific, fast-evolving GATA-like gene regulates zygotic gene activation to promote endoderm specification and pattern formation in the Theridiidae spider. BMC Biol. 2022;20(1):223. Epub 20221006.

55. Ranganayakulu G, Zhao B, Dokidis A, Molkentin JD, Olson EN, Schulz RA. A series of mutations in the D-MEF2 transcription factor reveal multiple functions in larval and adult myogenesis in Drosophila. Dev Biol. 1995;171(1):169–81.

56. Crittenden JR, Skoulakis EMC, Goldstein ES, Davis RL. Drosophila mef2 is essential for normal mushroom body and wing development. Biol Open. 2018;7(9). Epub 20180907.

57. Spahn P, Huelsmann S, Rehorn KP, Mischke S, Mayer M, Casali A, et al. Multiple regulatory safeguards confine the expression of the GATA factor Serpent to the hemocyte primordium within the Drosophila mesoderm. Dev Biol. 2014;386(1):272–9. Epub 20131217.

58. de Velasco B, Mandal L, Mkrtchyan M, Hartenstein V. Subdivision and developmental fate of the head mesoderm in Drosophila melanogaster. Dev Genes Evol. 2006;216(1):39–51. Epub 20051025.

59. Holz A, Bossinger B, Strasser T, Janning W, Klapper R. The two origins of hemocytes in Drosophila. Development. 2003;130(20):4955–62. Epub 20030820.

60. Tepass U, Fessler LI, Aziz A, Hartenstein V. Embryonic origin of hemocytes and their relationship to cell death in Drosophila. Development. 1994;120(7):1829–37.

61. Konigsmann T, Turetzek N, Pechmann M, Prpic NM. Expression and function of the zinc finger transcription factor Sp6-9 in the spider Parasteatoda tepidariorum. Dev Genes Evol. 2017;227(6):389–400. Epub 20171107.

62. Setton EVW, Sharma PP. Cooption of an appendage-patterning gene cassette in the head segmentation of arachnids. Proc Natl Acad Sci U S A. 2018;115(15):E3491–E500. Epub 20180326.

63. Schacht MI, Schomburg C, Bucher G. six3 acts upstream of foxQ2 in labrum and neural development in the spider Parasteatoda tepidariorum. Dev Genes Evol. 2020;230(2):95–104. Epub 20200210.

64. Steinmetz PR, Urbach R, Posnien N, Eriksson J, Kostyuchenko RP, Brena C, et al. Six3 demarcates the anterior-most developing brain region in bilaterian animals. Evodevo. 2010;1(1):14. Epub 20101229.

65. Cai J, St Amand T, Yin H, Guo H, Li G, Zhang Y, et al. Expression and regulation of the chicken Nkx-6.2 homeobox gene suggest its possible involvement in the ventral neural patterning and cell fate specification. Dev Dyn. 1999;216(4-5):459–68.

66. Gomez-Skarmeta JL, Modolell J. Iroquois genes: genomic organization and function in vertebrate neural development. Curr Opin Genet Dev. 2002;12(4):403–8.

67. Curtiss J, Heilig JS. Establishment of Drosophila imaginal precursor cells is controlled by the Arrowhead gene. Development. 1995;121(11):3819–28.

68. Sagasti A, Hobert O, Troemel ER, Ruvkun G, Bargmann CI. Alternative olfactory neuron fates are specified by the LIM homeobox gene lim-4. Genes Dev. 1999;13(14):1794–806.

69. Leszczynski P, Smiech M, Parvanov E, Watanabe C, Mizutani KI, Taniguchi H. Emerging Roles of PRDM Factors in Stem Cells and Neuronal System: Cofactor Dependent Regulation of PRDM3/16 and FOG1/2 (Novel PRDM Factors). Cells. 2020;9(12). Epub 20201204.

70. Hohenauer T, Moore AW. The Prdm family: expanding roles in stem cells and development. Development. 2012;139(13):2267–82.

71. Janssen R, Pechmann M, Turetzek N. A chelicerate Wnt gene expression atlas: novel insights into the complexity of arthropod Wnt-patterning. Evodevo. 2021;12(1):12. Epub 20211109.

72. Janssen R, Le Gouar M, Pechmann M, Poulin F, Bolognesi R, Schwager EE, et al. Conservation, loss, and redeployment of Wnt ligands in protostomes: implications for understanding the evolution of segment formation. BMC Evol Biol. 2010;10:374. Epub 20101201.

73. Giraldez AJ, Copley RR, Cohen SM. HSPG modification by the secreted enzyme Notum shapes the Wingless morphogen gradient. Dev Cell. 2002;2(5):667–76.

74. Tabata T, Eaton S, Kornberg TB. The Drosophila hedgehog gene is expressed specifically in posterior compartment cells and is a target of engrailed regulation. Genes Dev. 1992;6(12B):2635–45.

75. Morata G, Lawrence PA. Control of compartment development by the engrailed gene in Drosophila. Nature. 1975;255(5510):614–7.

76. Janssen R, Damen WG. Diverged and conserved aspects of heart formation in a spider. Evol Dev. 2008;10(2):155–65.

77. Tingvall TO, Roos E, Engstrom Y. The GATA factor Serpent is required for the onset of the humoral immune response in Drosophila embryos. Proc Natl Acad Sci U S A. 2001;98(7):3884–8. Epub 20010306.

78. Jacobs CG, Spaink HP, van der Zee M. The extraembryonic serosa is a frontier epithelium providing the insect egg with a full-range innate immune response. Elife. 2014;3. Epub 20141209.

79. Ferguson EL. Conservation of dorsal-ventral patterning in arthropods and chordates. Curr Opin Genet Dev. 1996;6(4):424–31.

80. Pechmann M, Benton MA, Kenny NJ, Posnien N, Roth S. A novel role for Ets4 in axis specification and cell migration in the spider Parasteatoda tepidariorum. Elife. 2017;6. Epub 20170829.

81. Holley SA, Neul JL, Attisano L, Wrana JL, Sasai Y, O’Connor MB, et al. The Xenopus dorsalizing factor noggin ventralizes Drosophila embryos by preventing DPP from activating its receptor. Cell. 1996;86(4):607–17.

82. Hunnekuhl VS, Akam M. An anterior medial cell population with an apical-organ-like transcriptional profile that pioneers the central nervous system in the centipede Strigamia maritima. Dev Biol. 2014;396(1):136–49. Epub 20140926.

83. Boyan G, Williams L. Embryonic development of the insect central complex: insights from lineages in the grasshopper and Drosophila. Arthropod Struct Dev. 2011;40(4):334–48. Epub 20110305.

84. de Velasco B, Erclik T, Shy D, Sclafani J, Lipshitz H, McInnes R, et al. Specification and development of the pars intercerebralis and pars lateralis, neuroendocrine command centers in the Drosophila brain. Dev Biol. 2007;302(1):309–23. Epub 20060926.

85. Posnien N, Koniszewski ND, Hein HJ, Bucher G. Candidate gene screen in the red flour beetle Tribolium reveals six3 as ancient regulator of anterior median head and central complex development. PLoS Genet. 2011;7(12):e1002416. Epub 20111222.

86. Lilly B, O’Keefe DD, Thomas JB, Botas J. The LIM homeodomain protein dLim1 defines a subclass of neurons within the embryonic ventral nerve cord of Drosophila. Mech Dev. 1999;88(2):195–205.

87. Shawlot W, Behringer RR. Requirement for Lim1 in head-organizer function. Nature. 1995;374(6521):425–30.

88. Zhu J, Palliyil S, Ran C, Kumar JP. Drosophila Pax6 promotes development of the entire eye-antennal disc, thereby ensuring proper adult head formation. Proc Natl Acad Sci U S A. 2017;114(23):5846–53.

89. Stollewerk A, Schoppmeier M, Damen WG. Involvement of Notch and Delta genes in spider segmentation. Nature. 2003;423(6942):863–5.

90. Pueyo JI, Lanfear R, Couso JP. Ancestral Notch-mediated segmentation revealed in the cockroach Periplaneta americana. Proc Natl Acad Sci U S A. 2008;105(43):16614–9. Epub 20081016.

91. Chesebro JE, Pueyo JI, Couso JP. Interplay between a Wnt-dependent organiser and the Notch segmentation clock regulates posterior development in Periplaneta americana. Biol Open. 2013;2(2):227–37. Epub 20121219.

92. Brena C, Akam M. An analysis of segmentation dynamics throughout embryogenesis in the centipede Strigamia maritima. BMC Biol. 2013;11:112. Epub 20131129.

93. Green J, Akam M. Evolution of the pair rule gene network: Insights from a centipede. Dev Biol. 2013;382(1):235–45. Epub 20130626.

94. Clark E, Peel AD. Evidence for the temporal regulation of insect segmentation by a conserved sequence of transcription factors. Development. 2018. Epub 20180503.

95. Janssen R. Segment polarity gene expression in a myriapod reveals conserved and diverged aspects of early head patterning in arthropods. Dev Genes Evol. 2012;222(5):299–309. Epub 20120818.

96. Auman T, Vreede BMI, Weiss A, Hester SD, Williams TA, Nagy LM, et al. Dynamics of growth zone patterning in the milkweed bug Oncopeltus fasciatus. Development. 2017;144(10):1896–905. Epub 20170421.

97. Shimizu T, Bae YK, Muraoka O, Hibi M. Interaction of Wnt and caudal-related genes in zebrafish posterior body formation. Dev Biol. 2005;279(1):125–41.

98. Ma P, Yang X, Kong Q, Li C, Yang S, Li Y, et al. The ubiquitin ligase RNF220 enhances canonical Wnt signaling through USP7-mediated deubiquitination of beta-catenin. Mol Cell Biol. 2014;34(23):4355–66. Epub 20140929.

99. Prpic NM, Damen WG. A homolog of the hydrolase Notum is expressed during segmentation and appendage formation in the Central American hunting spider Cupiennius salei. Naturwissenschaften. 2005;92(5):246–9. Epub 20050416.

100. Kornberg T. Engrailed: a gene controlling compartment and segment formation in Drosophila. Proc Natl Acad Sci U S A. 1981;78(2):1095–9.

101. Lawrence PA, Casal J, Struhl G. hedgehog and engrailed: pattern formation and polarity in the Drosophila abdomen. Development. 1999;126(11):2431–9.

102. Feitosa NM, Zhang J, Carney TJ, Metzger M, Korzh V, Bloch W, et al. Hemicentin 2 and Fibulin 1 are required for epidermal-dermal junction formation and fin mesenchymal cell migration during zebrafish development. Dev Biol. 2012;369(2):235–48. Epub 20120706.

103. Rescan PY, Ralliere C. A Sox5 gene is expressed in the myogenic lineage during trout embryonic development. Int J Dev Biol. 2010;54(5):913–8.

104. Urbano JM, Dominguez-Gimenez P, Estrada B, Martin-Bermudo MD. PS integrins and laminins: key regulators of cell migration during Drosophila embryogenesis. PLoS One. 2011;6(9):e23893. Epub 20110916.

105. Liang D, Wang X, Mittal A, Dhiman S, Hou SY, Degenhardt K, et al. Mesodermal expression of integrin alpha5beta1 regulates neural crest development and cardiovascular morphogenesis. Dev Biol. 2014;395(2):232–44. Epub 20140919.

106. Domsch K, Schroder J, Janeschik M, Schaub C, Lohmann I. The Hox Transcription Factor Ubx Ensures Somatic Myogenesis by Suppressing the Mesodermal Master Regulator Twist. Cell Rep. 2021;34(1):108577.

107. Posnien N, Zeng V, Schwager EE, Pechmann M, Hilbrant M, Keefe JD, et al. A comprehensive reference transcriptome resource for the common house spider Parasteatoda tepidariorum. PLoS One. 2014;9(8):e104885. Epub 20140813.

108. Bolger AM, Lohse M, Usadel B. Trimmomatic: a flexible trimmer for Illumina sequence data. Bioinformatics. 2014;30(15):2114–20. Epub 2014/04/04.

109. Dobin A, Davis CA, Schlesinger F, Drenkow J, Zaleski C, Jha S, et al. STAR: ultrafast universal RNA-seq aligner. Bioinformatics. 2013;29(1):15–21. Epub 2012/10/30.

110. Bruna T, Hoff KJ, Lomsadze A, Stanke M, Borodovsky M. BRAKER2: automatic eukaryotic genome annotation with GeneMark-EP+ and AUGUSTUS supported by a protein database. NAR Genom Bioinform. 2021;3(1):lqaa108. Epub 2021/02/13.

111. Buchfink B, Reuter K, Drost HG. Sensitive protein alignments at tree-of-life scale using DIAMOND. Nat Methods. 2021;18(4):366–8. Epub 20210407.

112. Buchfink B, Xie C, Huson DH. Fast and sensitive protein alignment using DIAMOND. Nat Methods. 2015;12(1):59–60. Epub 20141117.

113. Bankevich A, Nurk S, Antipov D, Gurevich AA, Dvorkin M, Kulikov AS, et al. SPAdes: a new genome assembly algorithm and its applications to single-cell sequencing. J Comput Biol. 2012;19(5):455–77. Epub 20120416.

114. Dierckxsens N, Mardulyn P, Smits G. NOVOPlasty: de novo assembly of organelle genomes from whole genome data. Nucleic Acids Res. 2017;45(4):e18.

115. Donath A, Juhling F, Al-Arab M, Bernhart SH, Reinhardt F, Stadler PF, et al. Improved annotation of protein-coding genes boundaries in metazoan mitochondrial genomes. Nucleic Acids Res. 2019;47(20):10543–52.

116. Martin M. Cutadapt removes adapter sequences from high-throughput sequencing reads. EMBnetjournal. 2011;17:10–2.

117. Macosko EZ, Basu A, Satija R, Nemesh J, Shekhar K, Goldman M, et al. Highly Parallel Genome-wide Expression Profiling of Individual Cells Using Nanoliter Droplets. Cell. 2015;161(5):1202–14.

118. McGinnis CS, Murrow LM, Gartner ZJ. DoubletFinder: Doublet Detection in Single-Cell RNA Sequencing Data Using Artificial Nearest Neighbors. Cell Syst. 2019;8(4):329–37 e4. Epub 20190403.

119. Patterson-Cross RB, Levine AJ, Menon V. Selecting single cell clustering parameter values using subsampling-based robustness metrics. BMC Bioinformatics. 2021;22(1):39. Epub 20210201.

120. Zappia L, Oshlack A. Clustering trees: a visualization for evaluating clusterings at multiple resolutions. Gigascience. 2018;7(7).

121. Yu G. Using ggtree to Visualize Data on Tree-Like Structures. Curr Protoc Bioinformatics. 2020;69(1):e96.

122. Prpic NM, Schoppmeier M, Damen WG. Whole-mount in situ hybridization of spider embryos. CSH Protoc. 2008;2008:pdb prot5068. Epub 20081001.

