## Supplementary text, tables and figures for "An atlas of spider development at single-cell resolution provides new insights into arthropod embryogenesis"

#### Split-pool ligation-based transcriptome sequencing (SPLiT-seq)

The SPLiT-seq protocol was performed as previously described with the following modifications [1].

##### Round 1 of Barcoding: Reverse Transcription

We prepared a 96-well plate with 8 μl/well of Round 1 barcodes from WD-1 (Working Dilution plate 1). With the plate on ice, we added 8 μl/well of the following RT mix: 4 μl of 5x Maxima RT Buffer (Thermo Scientific), 0.375 μl of Superase-In RNAse inhibitor (20 U/μl, Invitrogen), 1 μl of 10 mM/each dNTPs (NEB), 0.625 μl of nuclease-free water, and 2 μl of Maxima H Minus RT (200 U/µl, Thermo Scientific). Then, we added 8 μl/well of dissociated cells previously diluted in 0.5x PBS, at a concentration of 625 events/μl (5000 total events/well). The different embryonic stages were allocated to different columns of the plate, so each stage was labelled with a known-specific set of barcodes in this first round. The plate was incubated in a thermocycler for 35 min at 50ºC, and immediately place on ice afterwards. The individual reactions were pooled in a 15 ml Falcon tube, on ice. We added 9.6 μl of 10% Triton X-100 (~0.1% final concentration), mixed gently and centrifuged the cells at 1200 g for 5 min (4ºC). We removed the supernatant and resuspended de pellet in 2 ml of 1x NEB buffer 3.1 (NEB) with 20 μl of Superase-In RNase Inhibitor.

##### Round 2 of Barcoding: Ligation 1

The ligation mix was prepared with 500 μl of T4 Ligase Buffer 10x (NEB), 100 μl of T4 DNA ligase (400 U/μl, NEB), 100 μl of 1x PBS-1%BSA buffer and 1340 μl of nuclease-free water. The remaining steps were performed as previously described [1].

##### Round 3 of Barcoding: Ligation 2

For the second ligation, we added 150 μl of T4 DNA ligase (400 U/μl, NEB) to the pooled cells and mixed thoroughly by pipetting up and down. Then the Round 3 plate was filled with 55 μl/well of this cell-ligase mix. The remaining steps were performed as previously described [1].

##### Washing

After Round 3 of barcoding, we added 81 μl of 10% Triton-X 100 (~0.1% final concentration) to our pooled cells and centrifuged at 1200 g for 5 min (4ºC). The supernatant was carefully discarded, and cells were resuspended in 4.04 ml of washing buffer (4 ml of 1× PBS and 40 μl of 10% Triton X-100). Cells were centrifuged a second time, at 1200 g for 5 min (4ºC), and resuspended in 800 μl of 1x PBS-1% BSA buffer. The resuspended sample was transferred to a 1.5 ml Eppendorf tube and stored at -80ºC in 10% DMSO.

##### FACS

We FACS-sorted our cells in the middle of the SPLiT-seq protocol, after the barcoding rounds and before the cell lysis. To prepare the cells for the FACS, we thawed our sample on ice, added 4 μl of 10% Triton X-100 and centrifuged at 1200 g for 5 min (4ºC) to eliminate the DMSO. The supernatant was discarded and the pellet washed in 800 μl of 1x PBS-1% BSA buffer and, after adding another 4 μl of 10% Triton X-100, cells were centrifuged again in the same conditions. We finally resuspended the pellet in 800 μl of 1x PBS-1% BSA buffer and stained the cells with 1 μl of DRAQ5 (5 mM stock solution, Bioscience) and 2 μl of Concanavalin-A conjugated with AlexaFluor 488 (1 mg/ml stock solution, Invitrogen). The sample was kept in the dark, on ice, to stain for 45 minutes.

We sorted our cells in a BD FACS Aria III cell sorter (BD Biosciences) with a nozzle of 85 μm. We used the BD FACS Diva Software (BD Biosciences), setup in 4-Ways Purity Mode and moderate-pressure separation (45 Psi). We set DRAQ5-positive & Concanavalin-A-positive singlets to be sorted using the same gating strategy described in the *Flow cytometry* section. In total, we sorted two sub-libraries of ~15,000 cells each. Cells were collected in 1.5 ml eppendorf tubes using 50 μl of 2x lysis buffer as collection buffer, so cells were lysed as they were sorted. The sorting volume was adjusted to be approximately 50 μl per 15,000 cells, giving a final volume of 100 μl/tube and diluting the Lysis Buffer to a working concentration of 1x. The injection and collection chambers were kept at room temperature during the sorting run, which took approximately 1 hour.

##### Cell lysis

We added 10 μl of Proteinase K (20 mg/ml) to each sample and incubated for 2 h at 55ºC. After incubation, samples were stored at -80ºC.

##### Template Switch

The Template Switch mix was prepared using 44 μl of 5x Maxima RT Buffer (Thermo Scientific), 44 μl of 20% Ficoll PM 400 (Sigma Aldrich), 22 μl of 10 mM/each dNTPs (NEB), 5.5 μl of Superase-In RNAse inhibitor (20 U/μl, Invitrogen), 5.5 μl of *TSO primer (100 μM), 11 μl of Maxima H Minus RT (200 U/µl, Thermo Scientific), and 88 μl of nuclease-free water per sample. The Template Switch reaction was performed as previously described [1].

##### PCR Amplification

Samples were amplified for 5 cycles of PCR and, subsequently, for 10 cycles of qPCR. After amplification, samples were collected into 1.5 ml eppendorf tubes and frozen at -20ºC in the qPCR mix.

##### Size selection

Samples were thawed and purified in two rounds of SPRI size selection. We followed the Kapa Pure Beads (Roche) “Cleanup of Fragmented DNA in NGS Workflows” protocol with the following modifications: washing steps were performed with 750 µl of 85% ethanol, and cDNA was eluted in 20 µl of nuclease-free water at 37ºC for 10 min.

First we cleaned our samples with a 0.8x ratio of beads. After this, we added 80 µl of nuclease-free water to the elution (20 µl), to increase the sample volume up to 100 µl. Samples were then cleaned with a 0.7x ratio of beads and eluted in 20 µl of nuclease-free water.

Fragments distribution was assessed running a High Sensitivity DNA bioanalyzer (Agilent 2100). We also quantified the concentration of each sub-libraries using a Qubit dsDNA High Sensitivity Assay (Thermo Fisher).

##### Tagmentation

Samples were tagmented using the Nextera XT DNA Library Preparation Kit (Illumina). We diluted 1 ng of sample in 5 µl of nuclease-free water. Then, we prepared the tagmentation reaction with 5 µl of cDNA (1 ng), 10 µl of Tagment DNA Buffer (TD) and 5 µl of Amplicon Tagment Mix (ATM). Reactions were incubated in a pre-heated thermocycler at 55ºC for 5 min. Immediately after the incubation, we placed the samples on ice. To stop the tagmentation reaction, we added 5 µl of Neutralize Tagment Buffer (NT), mixed well, and incubated at room temperature for 5 min. Samples were kept on ice after incubation.

##### Round 4 of barcoding: PCR

For each sub-library, we prepared a separated PCR mix with 22 µl of tagmented sample, 15 µl of Nextera PCR Master Mix (Nextera XT DNA Library Preparation Kit), 1 µl of P5_oligo (10 µM) and 1 µl of Round 4 barcode (10 µM). We used a different Round 4 barcode per sub-library. In particular, we used the Round4_3 and Round4_4 oligos. The PCR reaction ran as follows: 72ºC (3 min); 95ºC (30 s); 12 cycles of 95ºC (10 s), 55ºC (30 s) and 72ºC (30 s); and 72ºC (5 min). Oligo sequences are provided in supplement of ACME methods [1]. After the PCR, samples were purified with two rounds of SPRI size selection (0.7x and 0.6x, subsequently) as described in the section *Size selection.* Libraries’ fragments distribution was assessed running a High Sensitivity DNA bioanalyzer (Agilent 2100) and final concentration was quantified using a Qubit dsDNA High Sensitivity Assay (Thermo Fisher).

#### Gene cloning and expression analysis

For gene expression characterisation in *Parasteatoda tepidariorum* embryos we performed colorimetric in situ hybridisation (ISH) [2], fastred [3] and double fluorescent ISH [4] as previously described with minor modifications.

##### RNA extraction and cDNA synthesis

*P. tepidariorum* embryos of stages 5-9 were pooled and total RNA was extracted using QiAzol (Qiagen) reagent, following the manufacturer’s guidelines, and stored at -80°C. Total RNA was used in cDNA synthesis using the QuantiTect reverse transcription kit (Qiagen), according to the manufacturer’s guidelines.

##### Primer design and probe synthesis

Primer3web ([http://primer3.ut.ee](http://primer3.ut.ee/)) was used to design gene-specific primers, and T7 linker sequences were added to the 5’ end of the forward (GGCCGCGG) and reverse (CCCGGGGC) primers. A complete list of primers can be found in Sup Table 2. Probe synthesis required two rounds of standard PCR using OneTaq 2x Master Mix (New England Biolabs). The first PCR used the gene-specific primers, the resulting purified PCR product was used as a template for the second PCR that used the gene-specific forward primer and 3’ T7 universal reverse primer (S1 Table). PCR products were run on a 1% agarose gel and purified using the NucleoSpin Gel and PCR Clean-up kit (Macherey-Nagel). DIG/Fluorescein labelled RNA probes were synthesised using T3 or T7 polymerase (Roche) with either DIG or Fluorescein RNA Labelling Mix (Roche), according to the manufacturer's guidelines.

##### In situ hybridization

After dechorionation with bleach, embryos were fixed overnight at room temperature (RT) in a two-phase solution of heptane and 37% formaldehyde in PBS (2:1:1). After fixation, embryos were washed with 100% methanol, rocked for ≥30 minutes at RT, and stored at -20°C. The vitelline membranes of fixed embryos were removed with Dumount 5 forceps in methanol.

For colorimetric ISH, minor modifications were made to the whole-mount ISH protocol [2]. Fixed embryos were gradually moved from methanol to PBS-T, the protein Proteinase K Digestion (Steps 4-8) was replaced by two 10-minute washes in PBS-Tween-20 (0.02%) (PBS-T), at step 18 embryos were incubated for 30 minutes, and at step 19 embryos were incubated for 2 hours. When required, embryos were ethanol treated to decrease background: embryos were washed for 5 minutes in 50% ethanol in PBS-T, washed in 100% ethanol until background had decreased, washed for 5 minutes in 50% ethanol in PBS-T, and finally washed twice with PBS-T.

For double fluorescent ISH, the protocol described was followed with some modifications [4]. The post-fixation, pre-hybridisation, hybridisation and probe removal steps follow steps 1-18 in the ISH protocol subject to the modifications previously stated [2]. Embryos were incubated with 1:2000 AP-conjugated anti-DIG (Roche) and 1:2000 POD-conjugated anti-FITC antibody (Roche) for 2 hours at RT and washed at least four times with PBS-T for 15 minutes each. Tyramide biotin amplification (TSA Plus Biotin Kit, Perkin–Elmer) was performed for 5-10 minutes, followed by a 90-minute incubation in 1:500 streptavidin Alexa Fluor 647 conjugate (ThermoFisher Scientific) to visualise the POD signal. The AP signal was visualised by a Fast Red reaction (Kem En Tec Diagnostics).

Embryos were counterstained with DAPI (Roche) in PBS-T (1:2000) for 20 mins at room temperature and stored in PBS-T at 4 °C. Embryos were flat-mounted in PBS-T on poly-L-lysine-coated coverslips and mounted with 80% glycerol or imaged wholemount in PBS-T. Colorimetric ISH and double fluorescent ISH were imaged using the Zeiss AxioZoom V16 or Zeiss LSM800 Confocal with Airyscan, respectively. Images were processed using FIJI software and Adobe Photoshop CS6.

### References

1.	Garcia-Castro H, Kenny NJ, Iglesias M, Alvarez-Campos P, Mason V, Elek A, et al. ACME dissociation: a versatile cell fixation-dissociation method for single-cell transcriptomics. Genome Biol. 2021;22(1):89. Epub 20210408.

2.	Prpic NM, Schoppmeier M, Damen WG. Whole-mount in situ hybridization of spider embryos. CSH Protoc. 2008;2008:pdb prot5068. Epub 20081001.

3.	Janeschik M, Schacht MI, Platten F, Turetzek N. It takes Two: Discovery of Spider Pax2 Duplicates Indicates Prominent Role in Chelicerate Central Nervous System, Eye, as Well as External Sense Organ Precursor Formation and Diversification After Neo- and Subfunctionalization Frontiers in Ecology and Evolution. 2022;10.

4.	Baudouin-Gonzalez L, Schoenauer A, Harper A, Blakeley G, Seiter M, Arif S, et al. The Evolution of Sox Gene Repertoires and Regulation of Segmentation in Arachnids. Mol Biol Evol. 2021;38(8):3153-69.

| Sup Table 1: Sequencing and cell processing metrics | | | | | | | | | |
| --- | --- | --- | --- | --- | --- | --- | --- | --- | --- |
| Per library | Raw read pairs | Trimmed and with barcode and paired reads used in mapping | Mapped reads (%) | Per stage | Dropseq processed reads (MG 100; MQ 0) | Number of cells after Seurat filtering and doublet removal | Total reads | Median UMI/cell | Median gene/cell |
| Library1 | 214,176,509 | 142375630 | 116897486 (82) | St7_1 | 3359678 | 1869 | 2790201 | 1360 | 612 |
|  |  |  |  | St8_1 | 3790082 | 1955 | 3263116 | 1724 | 657 |
|  |  |  |  | St9_1 | 7764729 | 3673 | 5371485 | 1299 | 534 |
| Library2 | 267,564,718 | 176303115 | 144913297 (82) | St7_2 | 5538607 | 2955 | 4923719 | 1538 | 717 |
|  |  |  |  | St8_2 | 6053367 | 2878 | 5343018 | 1724 | 756 |
|  |  |  |  | St9_2 | 10404026 | 5186 | 8019408 | 1369 | 587 |

Sup Table 2: Primers used for synthesis of in situ hybridisation probes

| Gene ID | Gene Name | Forward primer | Reverse primer | Size |
| --- | --- | --- | --- | --- |
| aug3.g11431 | *hamlet* | TTGCGACCCATTCCTTCGAA | CTCGTTGGCCTTGGGAGAAT | 895 |
| g29950 | *Hox3a* | TGGCACATTCCATCATCACA | TTGTTTGCTGTTGGGTGACG | 837 |
| g12191 | *lim1a* | AGTGAGAGAGACTCCAGCGT | ACCCGAAAAATTGGGTCTGC | 601 |
| g12868 | *Pax6.2* | AGACAGGCTCGATCAAACCC | TGGCACTAGAATCACTACCGT | 460 |
| g12873 | *Pax6.1* | GCTGCATCGTCTCTGTTCAG | GGTGTTGATGCCTCCGTTAC | 742 |
| g18856 | *SoxE2* | GACGATGACCTGGACAATGC | GCGACGGGTGATCTTCTCTA | 688 |
| aug3.g3745 | *Tbx3* | ACCACCTGAAGACGGAAGAG | GTCTTTTGGCGACGGTGAAT | 703 |
| aug3.g3790 | *omb* | ATGACTCCAGCGACGATGAT | GGGGTGTCGATGGAGTTGTA | 744 |
| g1245 | *six3.1* | CCCTACCTGTTGCTCATCCA | GCGTATGATGATGGTGCCTG | 804 |
| g25543 | *six3.2* | ACAGTCCCATGTTCGTCCTT | GGGCTGCTACCGTGTAAATG | 703 |
| aug3.g27186 | *vsx* | AATCTGCTGCACGGGGAT | ACTACACGGGCAGTCTTCAG | 539 |
| g18090 | *tll* | CATTCTTCAACCACTGCCCC | TTTTCTCCAGCCTCACCAGT | 668 |
| aug3.g2323 | *BMPR* | TGCAGGGATGGTCAGGTAGA | GTTGAAGAGTTCGGGCGTTG | 1201 |
| g27229 | *noggin* | GATTCTGCTCGACAAAACTACAG | CAGTAATTCGTAATAGAGAGCAATG | 997 |
| g12201 | *Nkx-6.2* | TCAACACACCAGCAACAACA | GCAAATCTATCCCTCCATAGTGC | 421 |
| g28941 | *RGMA* | ACTGTGATCATCCGTCAGCA | CGGAAGTCAGTGAAAGAGCTG | 556 |
| g7463 | *LRR2* | ACAAGAAAGCGGTCTGAGGA | GAGCGGTAATGGAACGACAC | 778 |
| g11868 | *VitK-C* | TACGAAGACACAGAGACGCA | TCAGCACCCACTTATCGGAG | 766 |
| g1958 | *GPCPD* | CGTGAGATCAACGAGCCAATGAAATGC | CGAATCCCAAAAGAGCAGCGGTAGA | 800 |
| g14898 | *HSP* | TGATTTGGACTGTGTTGCTGGCTGA | TTGGGATCGCTATTGACACAGTGCA | 1099 |
| g6459 | *myo* | ACAATTCAGGCTTATGTCCGTGGCA | TTTCTTGCTGTTGATGGGGATCGCT | 1427 |
| g6385 | *CAD-like* | GTTAAGTCTGCCAAAGAGATCGG | GTTAAGTCTGCCAAAGAGATCGG | 1480 |
| g4744 | *GATA2* | AACGGTGGGGTTAGTAGTGG | CTTCATGGTGAGAGGTCGGT | 720 |
| aug3.g16893 | *ush* | GAACTGTCGCCGGTAATTCC | GACGGGCAATGTATGAGCTG | 927 |
| g29100 | *cytochrome p450* | GGGCATGCATTCGATCCTAC | CCTTCCAAAGACAAGCACCC | 876 |
| g5619 | *tbx20* | GATGTCCCACCTCACCAGAA | ATCAGGAGGAAGTTCACCGG | 926 |
| g4985 | *platelet glycoprotein* | CAAGCGCGCAAAATAAAGCA | ATTGACGAGCTTGAGGGGAA | 857 |
| g3542 | *Mef2.1* | CGCAACTCCCCCAATCCGC | AGGAGGATGTAGATGGGGCAAGTGA | 1200 |
| g24898 | *Mef2.2* | AGGTCTTATGTTTCTGTGAGGTTGCCC | ATTGTTGGCAGTGTTATGTCCCCCA | 1300 |
| g11873 | *hemocyanin A* | AACAACAAGCACGGCAAAGAACTCC | GTCGCTTCGTTTTTGTAAAGGCCGT | 1250 |
| g472 | *C-ets1* | TTCCGAATCAAGTGCCACCTCTCAC | GTTTGAATTCCCACCCGTCTCCAGT | 1179 |
| g17128 | *DNA-directed RNA pol* | CTCTTAAGAAAATCGAACGATGGC | GATCAGAAAGCTAGTTTCTTAAGGAG | 447 |
| g13621 | *hemocyanin B chain* | CAGTATGTATCCATCTCTACCAAG | CATCAATGAACCTGTGCCATCTG | 1060 |
| g22680 | *hemocyanin C chain* | CATGCATCAACAGATGTGTGCTC | CTTAATGCCTGGCAGAATGGAC | 1289 |
| g23098 | *Integrin-a* | ATCAACTGCCAACTGTGCTG | TGAGCCCATCTGATCTCCTG | 616 |
| g30344 | *Notch2* | CAGCCAACCCTGTATGAACG | TCATGTCCAGTCCAGCCATT | 663 |
| g13852 | - | CCAATATAATGCACCCACGCT | AAGCGGCTTGAACAGTAGGA | 719 |
| g12621 | - | TGGTAGAAGGTGCAGTACGG | GGTGTTCCTTGGGATTGGTG | 701 |
| g21739 | *awh* | CAGGTAACATTGGCCAATGAATAAG | CAGGTAACATTGGCCAATGAATAAG | 986 |
| g18008 | *netrin* | GTGTGTGGCTTGCAACTGTA | GCCAGTCGTCTTTCCATTCC | 728 |
| g14495 | *latrophilin C* | AAATGGTATTCTGCCGTCGC | GGACGGTCGCAGCAAATATT | 660 |
| g15398 | - | CTAGTCGCTGTTCTTGTCGC | CCACCAGTTGATCCTTGCAG | 677 |
| aug3.g23531 | *AP2* | GGCCGCGGATTTCCCACCTCCCTACAGC | CCCGGGGCTACTTGTCGGGCAGGGAATT | 748 |
| g30822 | - | GGCCGCGGCCCGTTGGAGAAAAGCAACT | CCCGGGGCTACTCATGCATCTGACCGCT | 710 |
| g27156 | *RNF220* | CCCCAATCCACATTCTACGG | TGAGATCGTTCAGTTTGTCCA | 601 |
| g3028 | *DSPP* | GTTCCGACTCTAGTACGCCA | TGTGACAAAAGAGGCATGCC | 727 |
| g8835 | *big brother* | CGTGTAGTGCCCGATCAAGA | GCTCTTACCTCCACTTCTGC | 500 |
| g22446 | *band4.1* | TGTGATGGGAGAAGACAGCA | GGATATCTACGGGAGGGCAG | 708 |
| g17362 | *notum* | CCGGCAGGCTATTACATTCG | GATTTCGACACTGGAGCACC | 749 |
| g10589 | *gooseberry* | AGACCTGGTGTGATTGGAGG | GGATTGACGCATGGTTCAGG | 737 |
| g1871 | *hemicentin* | TCGTTCTGTCAGGTGGTTGA | ATTCTCTCGACGACTGGGTC | 952 |
| g29744 | *basonuclin* | AATCCCAAACTTCACACGCC | AAGACAGGTGGGGAGGAATG | 734 |
| g29290 | *Irx4* | AATGCCAGGAAAGTGACTCC | TGTTAGCTCTCACACCGACT | 835 |
| aug3.g13175 | *-* | TTGGGCATGCTCTGAAGGTA | ATGTCCTTCGCTGCTTTGTG | 767 |

**A**) rPCA

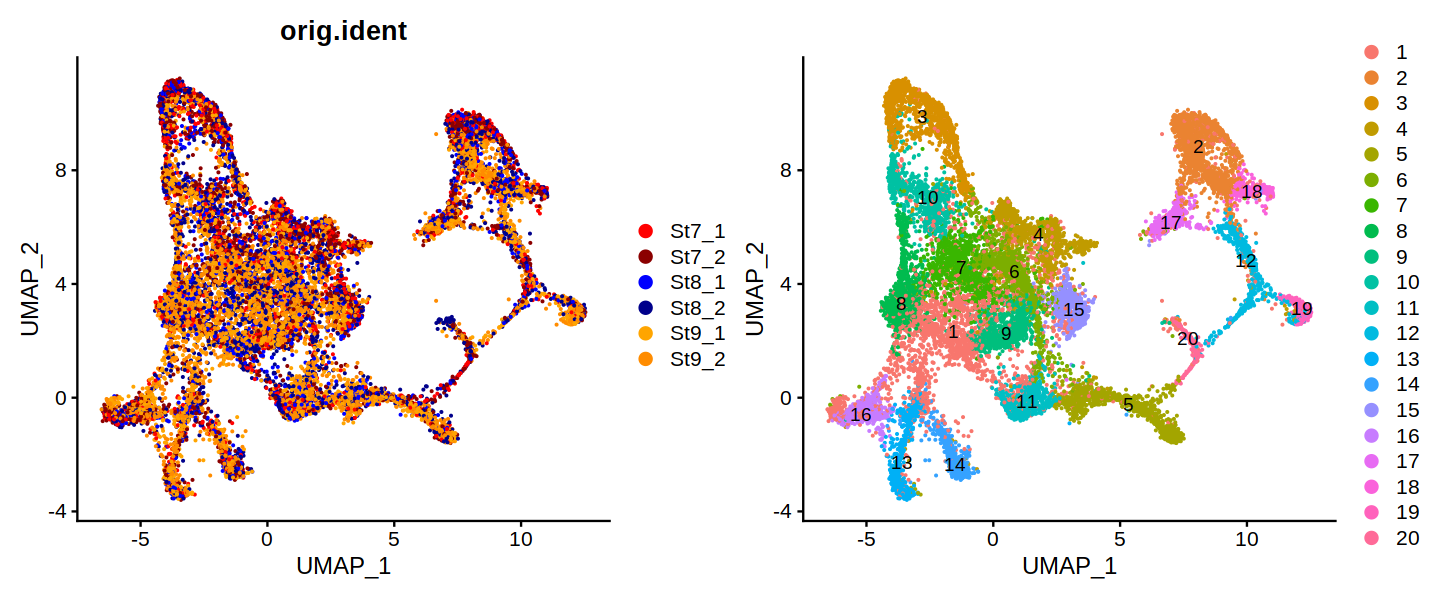

**B**) CCA

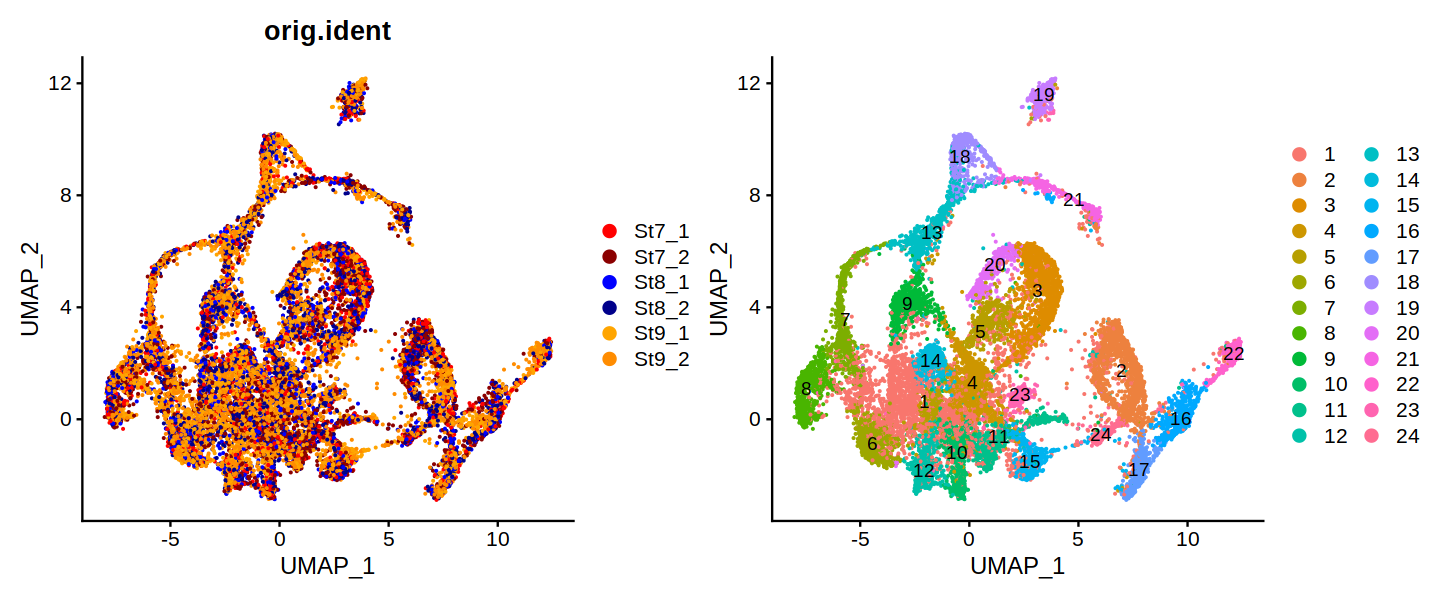

**C**) Harmony

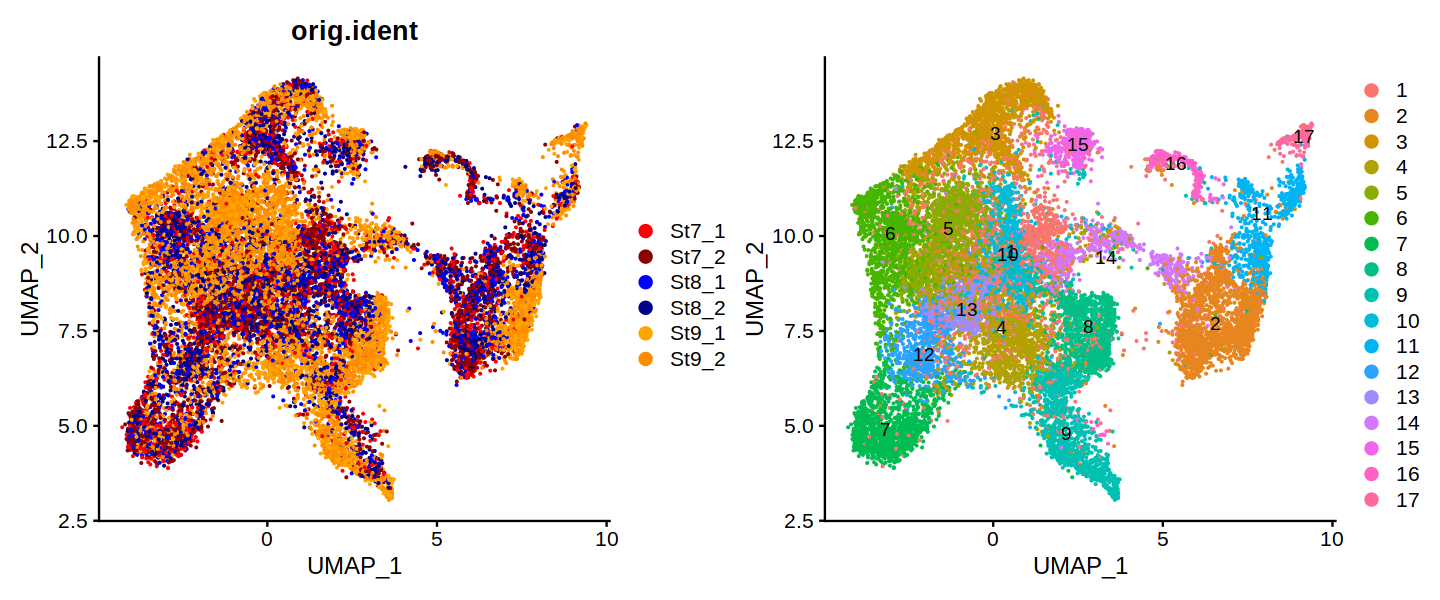

**Sup Fig 1: UMAPs showing sample integration methods.** (**A**) rPCA shows similar integration of stages as (**B**) CCA, whereas (**C**) Harmony maintains descriptively visable separation of stage 9.1 from stages 7 and 8.1.

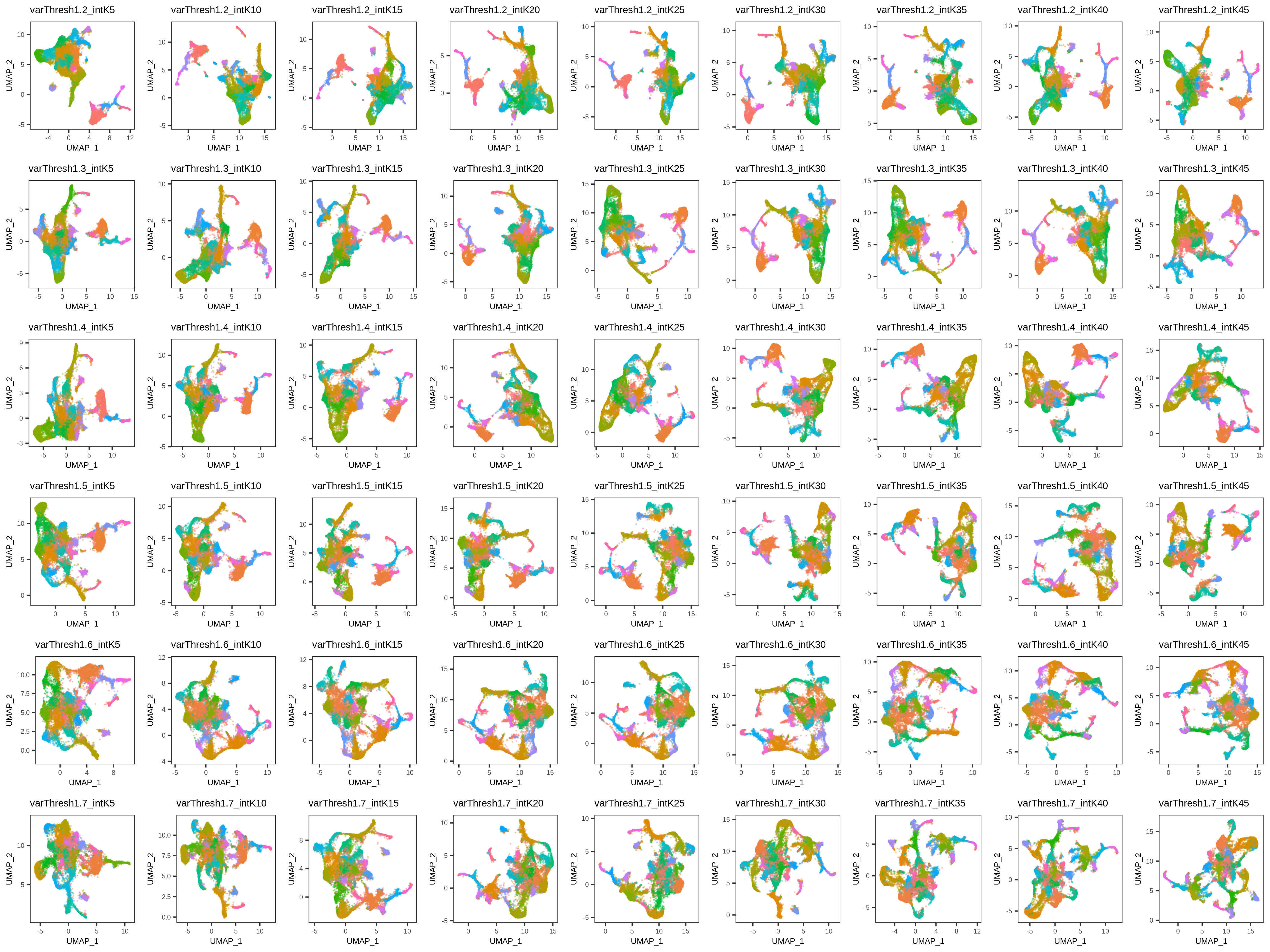

**Sup Fig 2: UMAPs of integration iterations.** Each row contains each iteration of highly variable gene (HVG) thresholds and each column is each iteration of the k.anchor value. All other parameters were fixed. These plots reveal that there are largely similar structures in the UMAPs between runs.

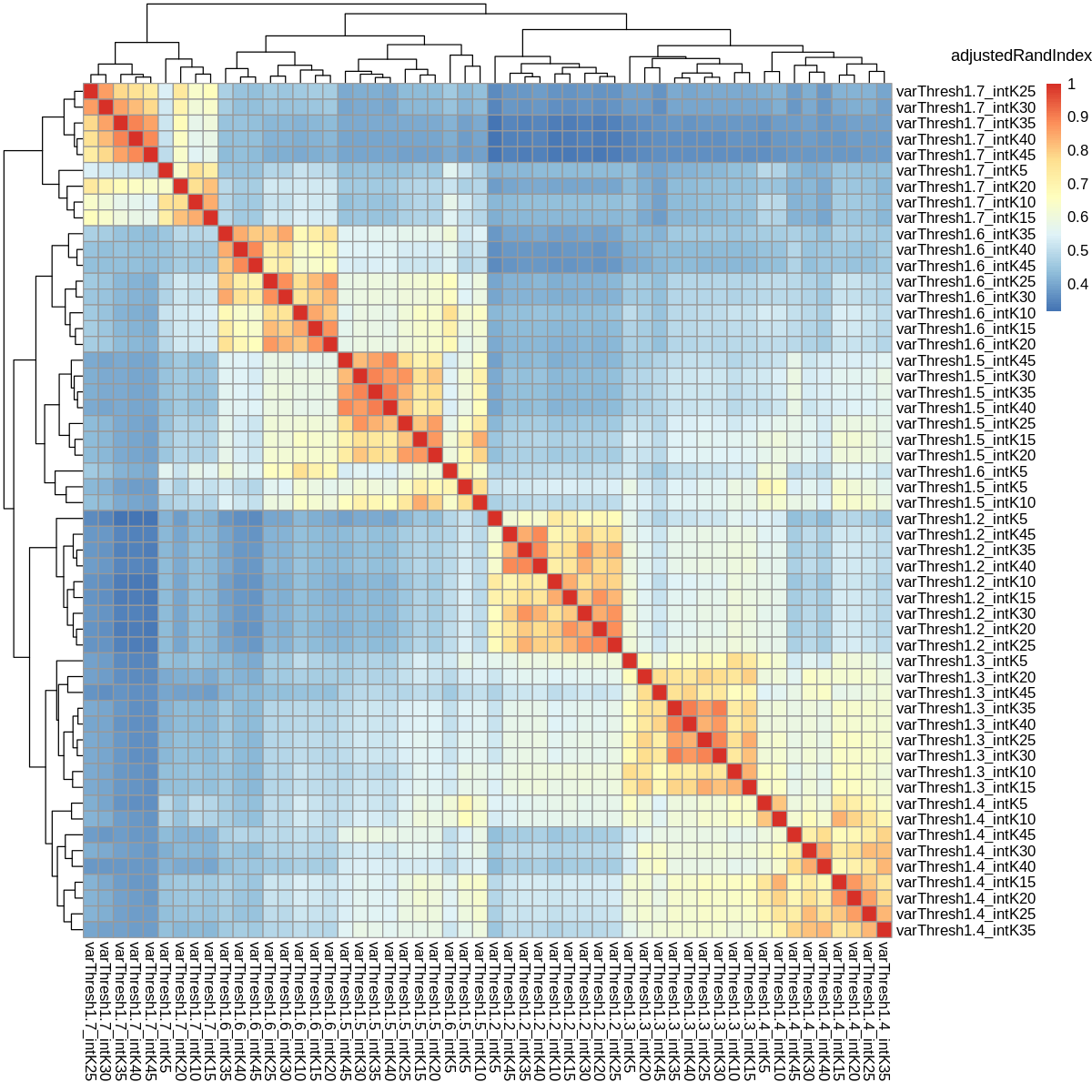

**Sup Fig 3: Heatmap of adjustRandIndex values comparing different integration iterations.** A quantitative assessment of clustering similarity between integration iterations showed that HVG threshold 1.2 is least similar to all others, whereas 1.3 and 1.4 are more comparable. k.anchors within each HVG threshold are more related. However, the combination of HVG threshold of 1.3 and anchors between 25-40 show the best overlap, suggestive that these might be reasonable parameter settings.

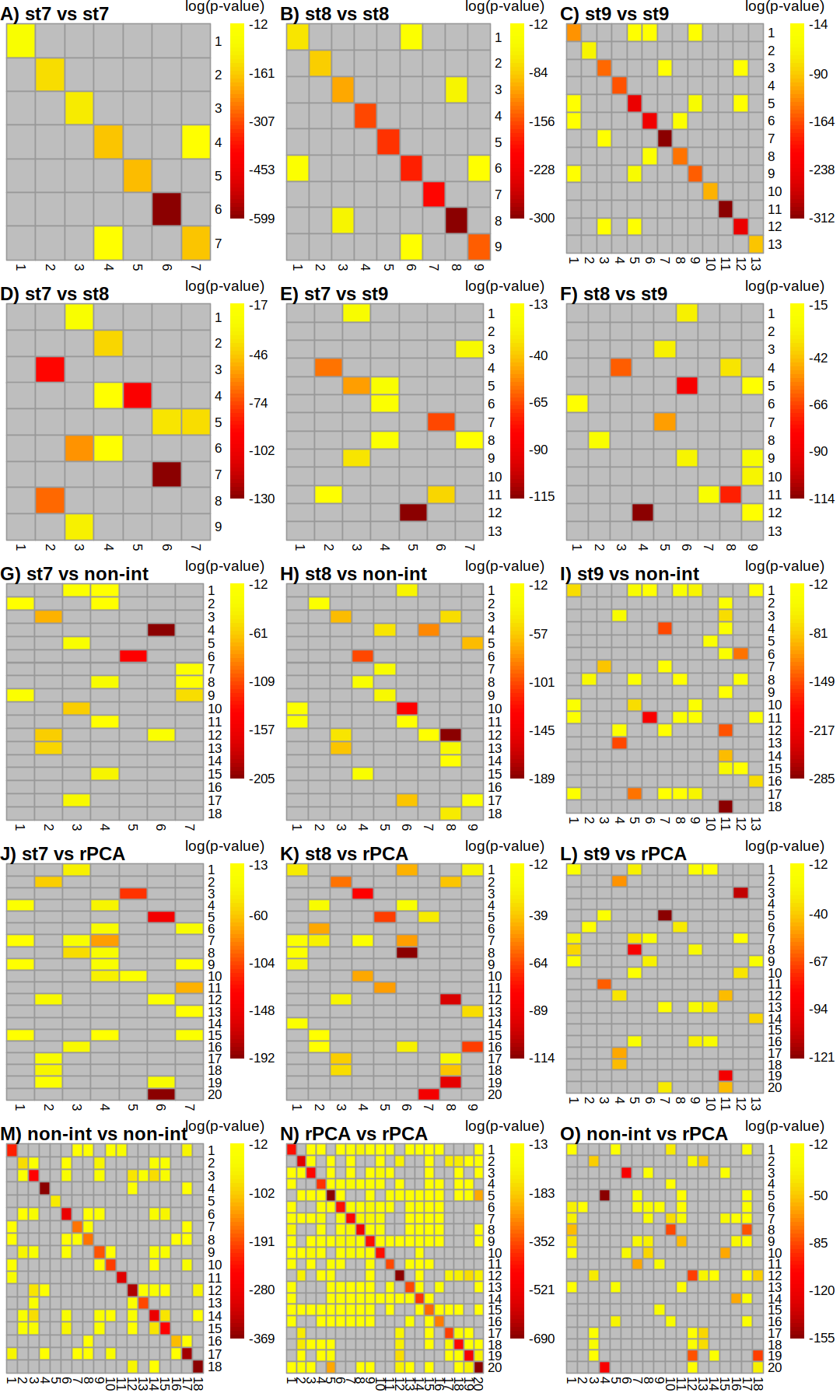

**Sup Fig 4: Hypergeometric distribution tests of cluster markers. (A** – **C)** stages compared to themselves show a lack of over-clustering and duplicates cell types. (**D** – **F)** stages compares to each other reveal similar cell cluster markers across stages. (**G** – **I)** stages compared to unintegrated merged data and (**J** – **L)** integrated data. (**M**) non-integrated self-comparison. (**N**) integrated self-comparison and (**O**) comparison between non-integrated and integrated.

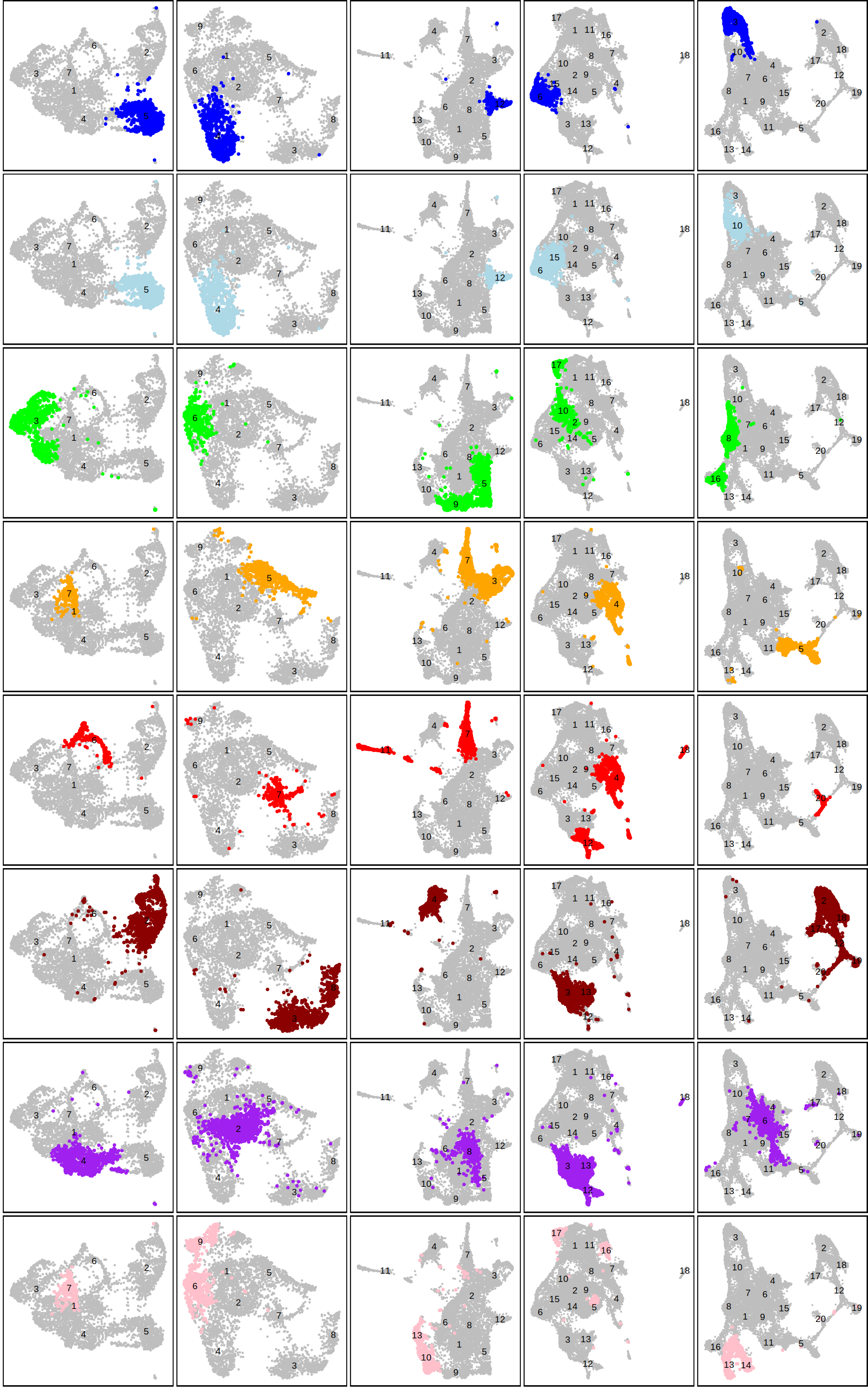

**Sup Fig 5: Matching clusters between stages, unintegrated and integrate data.** Using information from hypergeometric distribution test, similar cluster based of cluster markers between datasets were identified. These revealed clusters between datasets that were associated with SAZ (dark blue); SMZ (light blue); CNS & head (green), dorsal (orange); endoderm (red); mesoderm (brown); Posterior compartment (purple); head & precheliceral (pink).

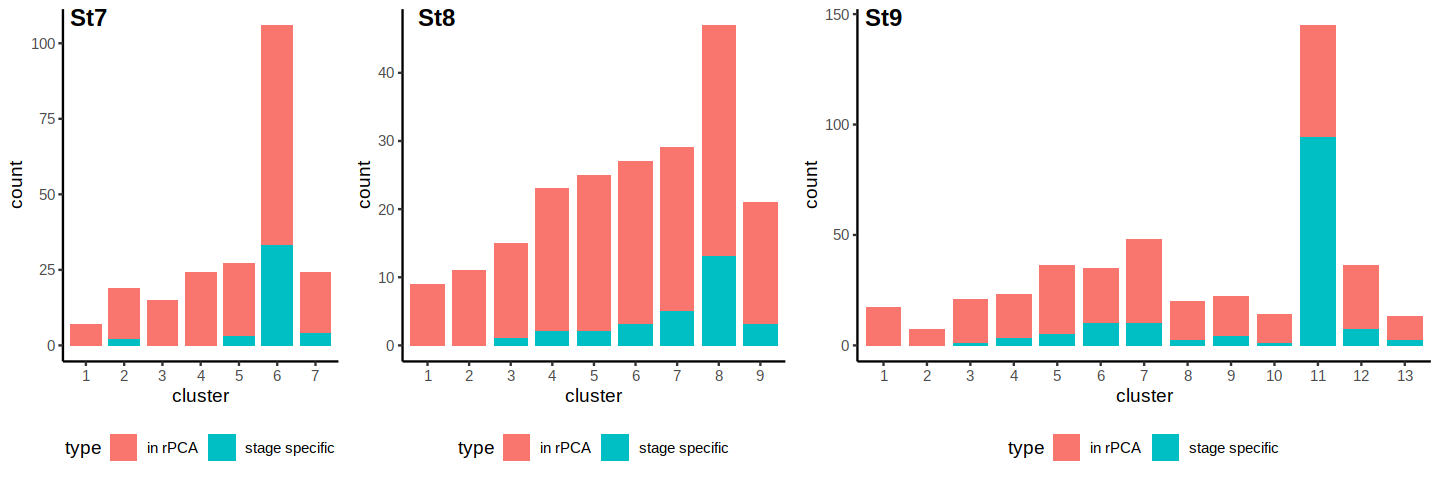

**Sup Fig 6: Numbers of stage specific cluster markers present/absent in integrated marker list.** Stage 7 and 8 markers were well captured in integrated markers. However, there were more stage 9 markers not found in the integrated marker list. This stage 9 cluster related very well with an integrated cluster (20), suggesting that while these markers were missed, the integrated clustering still captured reasonable information from these cells.

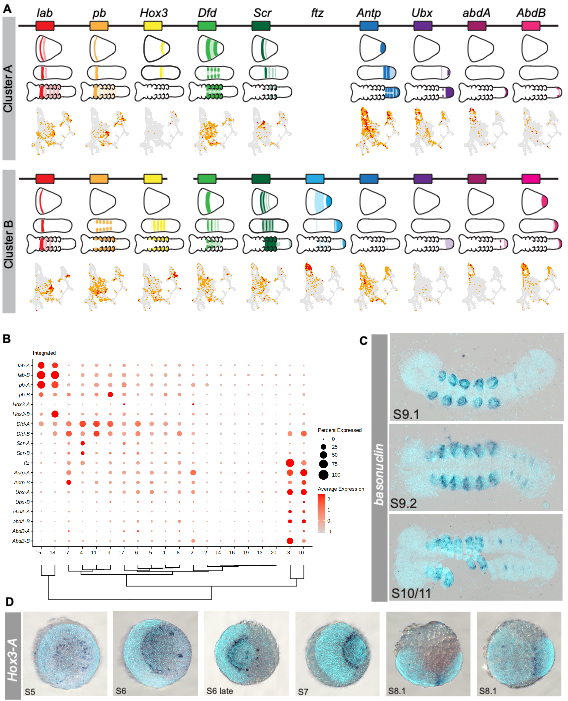

**Sup Fig 7: Hox expression in the scRNA-seq data.** (**A**) Schematics of Hox expression taken from in situ expression data in Schwager et al 2017 and UMAPs. (**B**) Dotplot for all Hox genes, not just hox markers. (**C**) Expression of cluster 9 marker *basonuclin* seen in the prosomal appendages and opisthosomal book lungs and spinnerets. (**D**) *Hox3-A* expression boarders the caudal lobe and prosoma, and later expressed in L2-L3.
